# The Lifespan Evolution of Individualized Neurophysiological Traits

**DOI:** 10.1101/2024.11.27.624077

**Authors:** Jason da Silva Castanheira, Alex I. Wiesman, Margot J. Taylor, Sylvain Baillet

## Abstract

How do neurophysiological traits that characterize individuals evolve across the lifespan? To address this question, we analyzed brief, task-free magnetoencephalographic recordings from over 1,000 individuals aged 4-89. We found that neurophysiological activity is significantly more similar between individuals in childhood than in adulthood, though periodic patterns of brain activity remain reliable markers of individuality across all ages. The cortical regions most critical for determining individuality shift across neurodevelopment and aging, with sensorimotor cortices becoming increasingly prominent in adulthood. These developmental changes in neurophysiology align closely with the expression of cortical genetic systems related to ion transport and neurotransmission, suggesting a growing influence of genetic factors on neurophysiological traits across the lifespan. Notably, this alignment peaks in late adolescence, a critical period when genetic factors significantly shape brain individuality. Overall, our findings highlight the role of sensorimotor regions in defining individual brain traits and reveal how genetic influences on these traits intensify with age. This study advances our understanding of the evolving biological foundations of inter-individual differences.

**Lay summary:** This study examines how brain activity reflects the development of individuality across a person’s life. Using magnetoencephalography to capture brief recordings of spontaneous brain activity, the researchers distinguished between over 1,000 individuals, spanning ages 4 to 89. They found that the brain regions most associated with individuality change with age: sensory and motor regions become increasingly distinctive in early adulthood, highlighting their role in shaping a person’s unique characteristics of brain activity. The study also revealed that changes in brain activity across different ages correspond to specific patterns of gene expression, shedding light on how genetics influence brain individuality. These findings deepen our understanding of the biological foundations of inter-individual differences and how it evolves over the lifespan.

## Introduction

The neurophysiological mechanisms underlying individuality in behavioural traits are a cornerstone of both basic and clinical neuroscience^1–8^. Beyond anatomical differences, functional neuroimaging studies have demonstrated that patterns of brain activity and connectivity reliably differentiate individuals^9–12^. These *neurophysiological traits*, referred to as “*brain-fingerprints*” in the literature, are associated with cognitive abilities^10,11,13,14^ and various clinical conditions, including neurological and psychiatric disorders^15–18^. Moreover, these traits are heritable and linked to cortical gene expression^19^. However, the evolution of these individual-specific neurophysiological features over the lifespan—amid extensive neurodevelopmental and aging-related brain changes—remains largely unexplored.

This study investigates how neurophysiological traits that differentiate individuals evolve across the lifespan, focusing on both developmental and aging trajectories. This question is particularly significant given the profound structural and functional transformations that occur from childhood through to older age. While behavioural traits and cognitive abilities change throughout life, it is unclear whether the underlying neurophysiological processes mirror these shifts or maintain continuity, reflecting a stable inter-individual differences.

A wealth of neuroimaging research has documented non-linear age-related changes in both brain structure and function, offering insights into lifespan development. Structurally, cortical thinning is a hallmark of aging, occurring progressively over time^20–23^. This thinning is particularly pronounced in higher-order association areas along the transmodal ends of the unimodal-to-transmodal functional gradient. Transmodal regions—associated with complex, integrative cognitive processes—show the earliest and most substantial reductions in cortical thickness^24,25^.

In terms of function, neurophysiological activity can be mapped using magnetoencephalography (MEG), which captures both periodic (oscillatory) and aperiodic (background) components of brain activity^26,27^. Periodic oscillations in different frequency bands (e.g., delta, theta, alpha, beta, and gamma) vary across individuals and systematically change with age. For instance, posterior dominant rhythms develop from slower (3–5 Hz) frequencies in early childhood to 6–7 Hz by 12 months, and then into the alpha rhythm (8–13 Hz) characteristic of adults^28–31^. This rhythm slows again in older age, reflecting functional decline^32–36^. Aperiodic components, which reflect the dynamic balance of excitatory and inhibitory processes^26,27,37–39^, also evolve across the lifespan and are associated with age-related declines in sensory and cognitive functions^36,40–44^.

These structural and functional transformations underscore the complex interplay between neurophysiological and anatomical changes over time. While behavioural traits and cognitive abilities evolve significantly, it remains an open question whether the neurophysiological processes differentiating individuals exhibit similar degrees of transformation or maintain stability. Prior research has shown that neurophysiological traits reliably differentiate young adults and can distinguish clinical populations, such as those with Parkinson’s disease, from age-matched healthy controls^9,15,16^. However, systematic exploration of how these the individual-specific traits change during key developmental stages—childhood, adolescence, and aging—is lacking.

This study addresses three key objectives: first, to determine whether the accuracy of individual differentiation based on neurophysiological traits changes with age; second, to explore how patterns of differentiation evolve across neurodevelopment and aging; and third, to identify the brain regions most critical for individual differentiation and understand their alignment with cortical gradients of functional organization and gene expression.

Previous studies of the development of neurophysiological traits have yielded mixed results. Some suggest that inter-individual differences in brain activity become more pronounced with age^17,45–47^, aligning with evidence of increasing genetic influence on cognition during development^48–50^. Other studies using functional magnetic resonance imaging (fMRI) have found that infants and children can be differentiated as accurately as adults, suggesting early stability in individual brain traits^51–55^. These discrepancies raise the question of whether neurophysiological traits become more individualized with age or whether distinct neurophysiology features differentiate individuals consistently throughout development.

Several factors may explain these mixed findings. Many studies rely on task-based data, where children and adults may employ different cognitive strategies, complicating differentiation. Additionally, fMRI—an indirect measure of brain activity sensitive to hemodynamic neural coupling—introduces potential age-dependent biases^56–58^. In contrast, MEG directly measures neurophysiological activity, offering a clearer perspective on how differentiation evolves.

Emerging evidence also indicates that the brain regions most characteristic of healthy individuals differ significantly from those in clinical populations, where disease-related disruptions alter neurophysiological patterns^15,16,59^. This suggests that features critical for differentiation may shift in response to neural alterations due to disease or the cumulative effects of aging and development. Supporting this, models trained on young adult data to predict behaviour from brain activity often fail to generalize when applied to older adults^60,61^, emphasizing the importance of age-specific patterns in studying individual differences.

In this study, we analyzed MEG-derived neurophysiological traits from a lifespan sample of 1,007 participants aged 4 to 89 years. We examined whether individuals could be differentiated across a wide age range and investigated how the topography of the most characteristic brain regions shifts with age. Our findings reveal that while periodic brain activity reliably differentiates individuals across the lifespan, the regions contributing most strongly to this differentiation change systematically with age, aligning with cortical gradients of functional organization and gene expression. These results reconcile previous inconsistencies in the developmental trajectory of brain individuality and offer new insights into how neurophysiological traits evolve throughout life.

## Results

We investigated three core aspects of neurophysiological traits across the lifespan. First, we evaluated whether individuals from different age groups, particularly children and older adults, could be accurately differentiated based on brief recordings of their neurophysiological brain activity. Next, we examined the ease with which we can differentiate individuals through neurodevelopment and aging. Finally, we assessed the spatial distribution of neurophysiological features that most strongly contribute to individual differentiation and analyzed how these features vary systematically with age and relate to established cortical gradients of functional organization and gene expression.

We used a neurophysiological trait differentiation method that replicates established brain-neurophysiological differentiation techniques^9–11^. This approach involved comparing each participant’s neurophysiological traits—derived from one segment of data—with traits from all other participants, including a separate segment from the same individual. Differentiation accuracy was determined based on whether the participant’s traits were more similar to each other (“self-similarity”) than to those of other individuals (“other-similarity”). This analysis quantified differentiation accuracy across age groups and provided insights into the stability and uniqueness of neurophysiological traits over time.

### Differentiation Accuracy Across Age Groups and Neurophysiological Components

We computed differentiation accuracy across four age cohorts from the SickKids dataset: i) children aged 4–12 years (N = 148), ii) adolescents aged 12–18 years (N = 57), iii) adults aged 18– 68 years (N = 196), and iv) all participants combined (N = 401).

Across all age groups, individuals were accurately differentiated with an overall accuracy of 93.3% (95% confidence interval [CI]: [92.2, 94.5]; Figure 1a). Differentiation accuracy for children was 90.2% (CI: [88.7, 92.5]), for adolescents 96.7% (CI: [94.1, 100.0]), and for adults 95.5% (CI: [94.3, 97.1]). These results were consistent when individual traits were derived from specific frequency bands and age-specific data subsets (Figures S1, S7).

**Figure 1:**
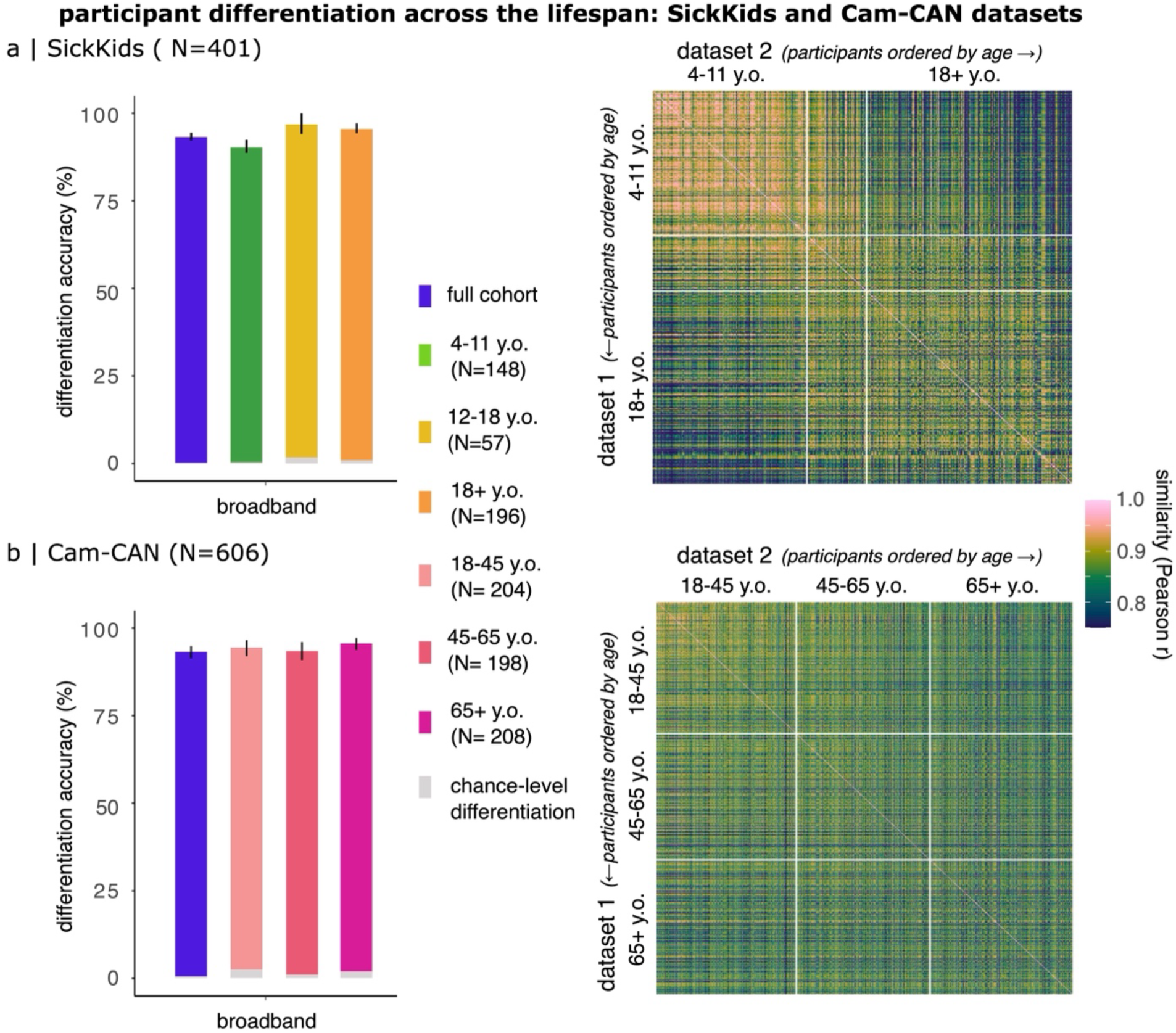
Age-related Variations in Neurophysiological Differentiation Accuracy. (a) Differentiation accuracy in the SickKids cohort (ages 4–68), using wide-band neurophysiological traits derived from 2-minute MEG recordings. Grey bars at the base of the plot represent chance-level differentiation based on empty-room MEG recordings collected on the same day as participants’ sessions. Error bars denote bootstrapped 95% confidence intervals. (b) Similarity matrix for the SickKids cohort, illustrating self-similarity (diagonal) and other-similarity (off-diagonal) across two distinct data segments collected from each participant. Participants are ordered by ascending age, with children (ages 4–11) exhibiting higher other-similarity values, making differentiation more challenging. (c) Differentiation accuracy in the Cam-CAN cohort (ages 18–89), using wide-band neurophysiological traits from 4-minute MEG recordings. Grey bars represent chance-level differentiation as in (a). Error bars denote bootstrapped 95% confidence intervals. (d) Similarity matrix for the Cam-CAN cohort, illustrating age-related changes in self- and other-similarity. Older adults exhibit higher distinctiveness in their neurophysiological traits, contributing to the modest positive relationship between differentiation and age (see Figure S2).

To investigate development trends, we assessed “differentiability”^9^, a measure of how easily individuals could be differentiated based on their neurophysiological traits. We observed a strong positive linear relationship between age and differentiability (r = 0.60, t(399) = 14.83, p < 0.001; Figure S2). This increase was primarily driven by greater other-similarity in children, which made differentiation more challenging in this group. In contrast, adult traits exhibited lower similarity with other adults, enabling more accurate differentiation (Figure 1a, right panel).

To further explore the contributions of periodic and aperiodic components to differentiation, we decomposed the power spectral density (PSD) of each MEG cortical source time series into these two components. These decomposed spectra were then used to define individual periodic and aperiodic spectral traits (see Methods).

When differentiation accuracy was computed using aperiodic traits, accuracy dropped to 78.8% overall (CI: [76.5, 80.9]), with 75.3% accuracy for children (CI: [72.9, 78.2]), 85.8% for adolescents (CI: [80.4, 92.2]), and 83.9% for adults (CI: [81.2, 86.9]; Figure S8). In contrast, differentiation based on periodic trats remained highly accurate across all age groups, achieving 96.5% overall accuracy (CI: [95.6, 97.5]), 96.8% for children (CI: [96.2, 97.7]), 97.4% for adolescents (CI: [96.1, 100.0]), and 96.6% for adults (CI: [95.5, 98.3]; Figure S9).

### Differentiation Accuracy and Stability in Older Adults

To extend our findings to older populations, we analyzed neurophysiological differentiation in an adult cohort (ages 18-89 years, N=606) from the Cambridge Center for Aging Neuroscience (Cam-CAN) dataset^62^. Participants were divided into three age groups: young adults (18-45 years, N = 204), middle-aged adults (45-65 years, N = 194), and older adults (65-89 years, N = 208).

For the entire Cam-CAN cohort, differentiation accuracy was 93.2% (CI: [91.3, 94.8]). Differentiation accuracy varied slightly across age groups: 94.4% (CI: [92.0, 96.6]) for young adults, 93.4% (CI: [90.8, 96.0]) for middle-aged adults, and 95.6% (CI: [93.7, 97.1]) for older adults (Figure 1b). These findings were robust across frequency bands and decomposition of periodic and aperiodic traits (Figure S1, S8, S9). Note, that the differentiation accuracy of adults in the SickKids dataset was similar to the young adults in the Cam-CAN cohort.

To assess whether the stability of individual traits changes with age, we analyzed variations in self-similarity across age groups. No significant linear relationship was observed between self-similarity and age (β = -4.50, SE = 2.45, 95% CI [-9.31, 0.31], r² = 0.006, p = 0.06). However, differentiability showed a modest positive relation with age (β = 5.84, SE = 1.12, 95% CI [3.62, 8.07], r² = 0.042, p < 0.001), explaining 4.2% of the total variance (Figure S2).

Finally, we conducted sensitivity analyses to rule out potential confounding effects from environmental or physiological artifacts in both cohorts’ data. These analyses (detailed in Figures S3—S5) demonstrated that artifacts did not significantly affect differentiation accuracy. Furthermore, anatomical similarity was not a significant contributor to the higher other-similarity observed in children’s neurophysiological traits (Figure S6).

### Age-Related Shifts in Cortical Regions Influential for Differentiation

Given the lower differentiation accuracy observed in children when using non-parameterized neurophysiological traits, we focused our analyses on periodic activity, which consistently demonstrated higher differentiation accuracy across all age groups.

Our primary goal was to identify the cortical regions most characteristic of individuals throughout the lifespan. We define “characteristic” regions as those where neurophysiological features most strongly contribute to individual differentiation. To quantify this, we used intraclass correlations (ICCs), which measure the consistency of neurophysiological features within individuals relative to variability across individuals^9,10,16^. Regions with higher ICC values were deemed more influential for individual differentiation. To capture developmental trajectories, we used a sliding window approach to compute ICC maps across different age groups (see Methods).

Our findings revealed systematic shifts in the cortical regions most distinctive for individual differentiation over the lifespan. In children, lateral parietal and superior temporal regions were the most characteristic of individual differences. By early adulthood (ages 20-30 years), caudal and peri-central cortical regions emerged as the most distinctive, consistent with previous studies^12,19^. Across all age groups, orbitofrontal regions consistently contributed the least to differentiation (Figure 2a and Figure S11).

**Figure 2:**
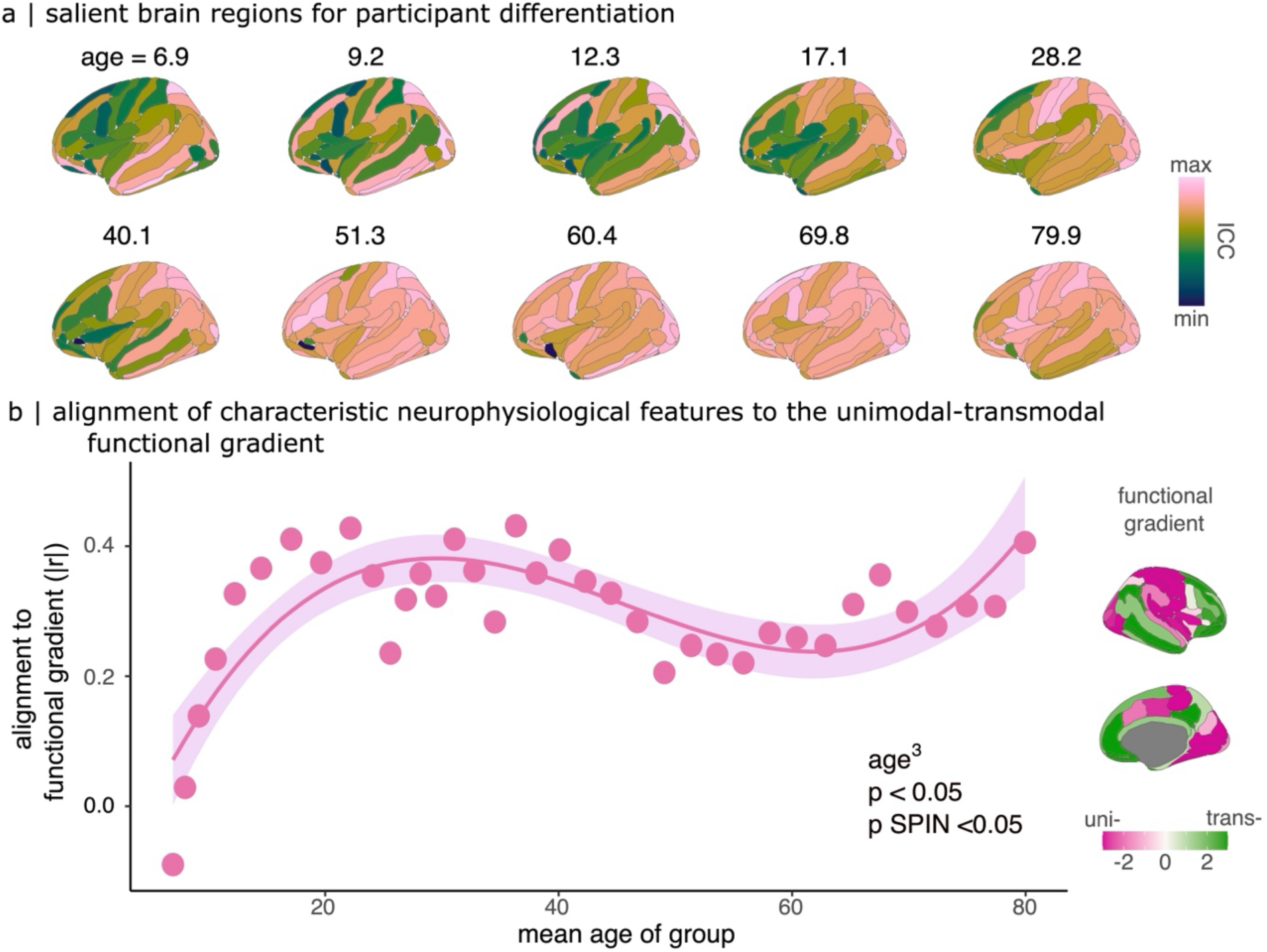
Sensorimotor Brain Regions Become Increasingly Characteristic for Individuals Across the Lifespan. (a) Topographic maps of intraclass correlation coefficients (ICCs) show the cortical regions that contribute most to individual neurophysiological differentiation across age groups (bins of ∼100 individuals per map). Higher ICC values indicate cortical areas that play a significant role in distinguishing individuals within each age group. Sensorimotor regions become increasingly prominent with age. (b) Scatter plot illustrating the alignment between characteristic cortical regions for participant differentiation (as shown in panel a) and the unimodal-to-transmodal functional gradient (inset on the right). The alignment exhibits a non-linear trajectory across the lifespan, with weak alignment in early childhood, progressively increasing into young adulthood, and peaking at 36.3 years old. This highlights the evolving contribution of cortical regions to individual differentiation with age.

Interestingly, these characteristic patterns of periodic neurophysiology also predicted cognitive performance. In the Cam-CAN sample, periodic features showed significant correlations with fluid intelligence test scores (Figure S10), suggesting that regions central to differentiation may also underlie variability in cognitive abilities.

We examined how these characteristic cortical regions align with the unimodal-to-transmodal functional gradient^63^, a primary axis of neurodevelopment that reflects transitions from primary sensory-motor areas to higher-order association cortices^24,25^. The alignment of characteristic regions with this gradient followed a non-linear trajectory across the lifespan (3^rd^ order polynomial: β = -0.46, SE = 0.07, 95% CI [-0.60, -0.31], p < 0.001, p_spin_ < 0.05; Table S6). Alignment was weakest in early childhood (6.9 years old, r = 0.09, p_spin_ = 0.26) and strongest in early adulthood (36.3 years old, r = -0.44, p_spin_ < 0.05; Figure 2b).

In children, we found no significant alignment between characteristic cortical regions and the unimodal-to-transmodal gradient (9.2 years old, r = -0.14, p_spin_ = 0.25; Figure 2b). This observation led us to hypothesize that the regions most characteristic of individual differences might instead align with the motor-to-visual gradient, a functional gradient crucial for early brain development^64^. Supporting this hypothesis, we observed a significant spatial alignment between the motor-to-visual gradient and the characteristic cortical regions in children (9.2 years old, r = 0.43, p_spin_ < 0.05; Figure S13).

### Alignment Between Neurophysiological Differentiation and Cortical Gene Expression Across the Lifespan

Previous research has shown that neurophysiological traits in adults align with a ventromedial-to-dorsolateral gene expression signature, involving genes associated with ion transport, synaptic functioning, and neurotransmitter release^19^. However, the evolution of this alignment across different developmental stages, particularly during critical periods of neurodevelopment, has remained unclear. We aimed to elucidate how the alignment between cortical neurophysiological traits and cortical gene expression changes across the lifespan (Figure 2a).

To investigate this relationship, we divided the gene-differentiation signature into two sets: a positive set related to participant differentiation, and a negative set associated with lower differentiation capabilities. The positive gene set primarily included processes related to ion transport, synaptic activity, and neurotransmitter release, while the negative set was linked to neurodevelopmental processes such as neurogenesis and cell morphogenesis^19^. This division allowed us to understand which gene expressions were related to neurophysiological differentiation across age groups (see Methods).

We first assessed whether the spatial organization of these gene sets remained stable across developmental stages, using data from the BrainSpan dataset^65^, which includes fetal through to adult data. Consistent with prior findings indicating that the most pronounced changes in cortical gene expression occur prenatally^66^, the spatial distribution of the positive gene signature showed remarkable stability from infancy through childhood, adolescence, and adulthood (spatial correlations > 0.71; Figure S14). In contrast, the negative gene signature exhibited weaker spatial consistency between childhood and adulthood (Figure S14). These results underscored the enduring relevance of the positive gene set for neurophysiological differentiation, prompting us to focus subsequent analyses on this gene group.

Next, we analyzed the alignment between cortical maps of neurophysiological differentiation and the positive gene signature. This alignment progressively strengthened with age, reaching its peak during late adolescence (17.1 years old; β = 0.73, SE = 0.11, 95% CI [0.51, 0.96], p < 0.001, p_spin_ < 0.01; Table S7, Figure 3a). This finding suggests that adolescence represents a critical period during which neurophysiological features most strongly align with genetic signatures related to ion transport and neurotransmission^19^, likely reflecting a pivotal stage in the fine-tuning of neural circuits.

**Figure 3:**
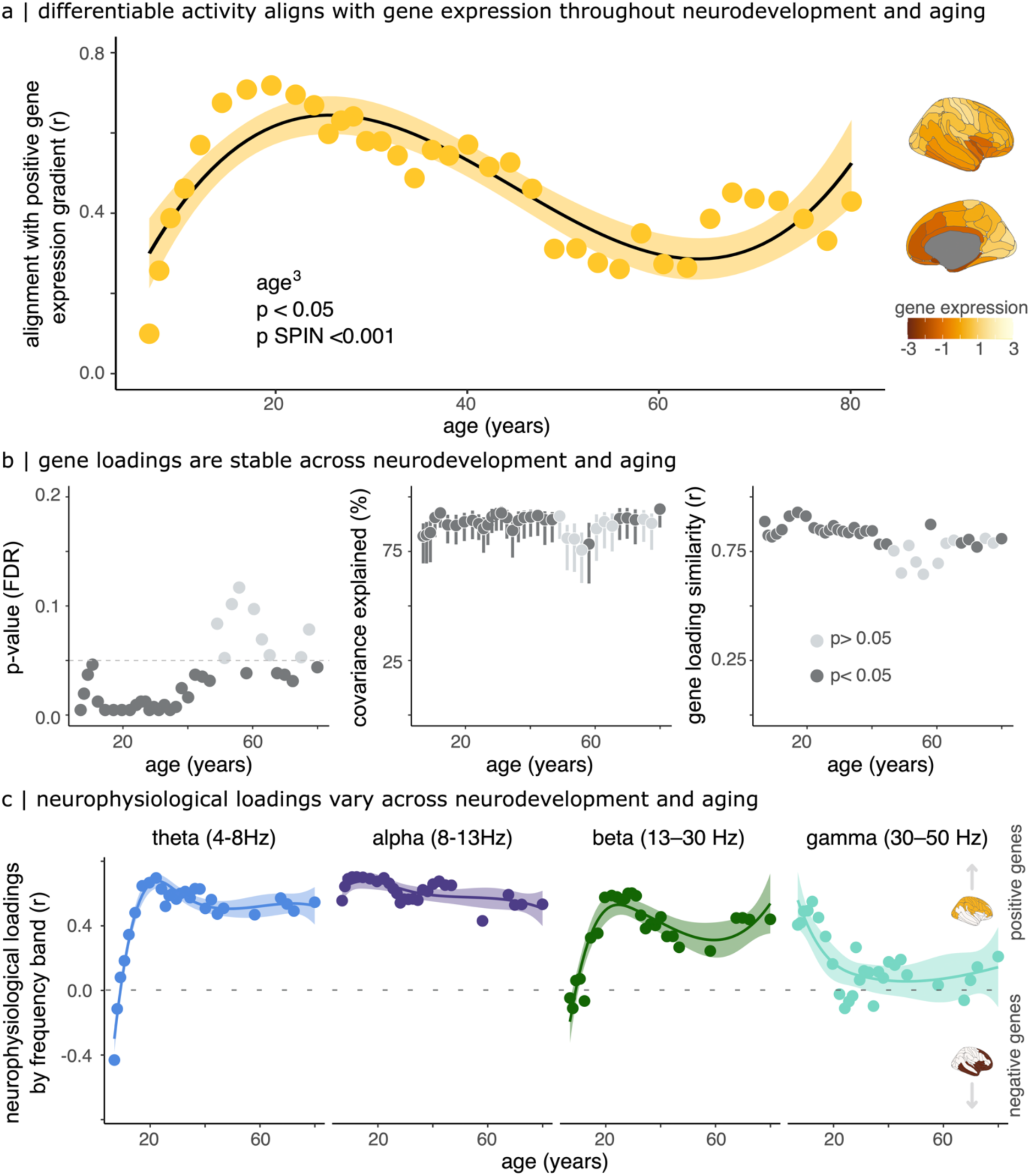
Lifespan Variations in the Alignment Between Gene Expression and Characteristic Patterns of Neurophysiological Activity. (a) Cortical alignment between the positive gene expression signature and differentiable brain regions: The right inset shows the spatial distribution of the positive gene expression signature across the cortex. The graph illustrates the alignment of this genetic signature with cortical regions most critical for neurophysiological differentiation across the lifespan. The alignment is weakest during early childhood (approximately 6.9 years old) and peaks in late adolescence (17.1 years old), emphasizing critical periods where genetic expression most influences neurophysiological individuality. (b) Multivariate PLS analysis results:

- Left: Significance (p-values) of latent gene differentiation components, revealing significant covariance between cortical gene expression and neurophysiological differentiation across the lifespan, persisting through middle age. Points above the dashed line did not survive correction for multiple comparisons (FDR) or spatial autocorrelation.
- Middle: Percent covariance explained by the latent components across different age groups. The variance explained remains relatively stable across age bins, suggesting a consistent gene-neurophysiology relationship throughout development and aging.
- Right: Gene loadings show the relative contribution of neurophysiological frequency bands (theta, alpha, beta, gamma) and genes to the covariance pattern. Strong consistency is observed across developmental stages, as well as with loadings from prior independent analyses, demonstrating the stability of gene contributions to neurophysiological differentiation over time. (c) Frequency band contributions across neurodevelopment: The graph depicts the relative contributions of neurophysiological frequency bands (theta [4-8 Hz), alpha [8-13 Hz), beta [13-30 Hz), gamma [30-50 Hz]) to gene-neurophysiology covariance. Positive neurophysiological loadings align with positive gene expression, while negative loadings align with negative gene expression (illustrated by topographies on the far right). Theta band activity peaks early in development, while beta and gamma band contributions increase later in life. Alpha band contributions remain stable across neurodevelopment and aging, underscoring their consistent role in brain individuality.

To gain deeper insights into how gene-neurophysiology alignment evolves, we conducted a Partial Least Squares (PLS) analysis to examine the covariance between neurophysiological features and cortical gene expression data from the Allen Human Brain Atlas^67^. The analysis revealed significant covariance between cortical gene expression patterns and neurophysiological differentiation across the lifespan, particularly up to middle age (Figure 3b, left). Interestingly, the percentage of covariance explained by these latent components remained relatively stable across age groups (Figure 3b, middle), indicating that the relation between gene expression and neurophysiological differentiation is preserved over time. Additionally, gene loadings showed strong consistency across developmental stages, with Pearson correlations > 0.75 between loadings from different age groups. This consistency suggests that the same gene signature drives neurophysiological differentiation throughout development (Figure 3b, right).

We further examined the contributions of specific neurophysiological frequency bands to the alignment with gene expression (Figure 3c). The relationship between periodic activity and gene expression varied significantly by frequency band and developmental stage. Theta band activity (4-8 Hz) was associated with negative gene expression during early development, and this association reversed later in life, potentially reflecting shifts in functional priorities as neurodevelopment progresses. Beta band activity (13-30 Hz) displayed a similar trend, showing a growing alignment with positive gene expression in adulthood, likely reflecting its critical role in sensorimotor functions and predictive coding. Gamma band activity (30-50 Hz) strongly aligned with positive gene expression during early development, but its contribution decreased with age, perhaps reflecting decreased cortical plasticity over time. Alpha band activity (8-13 Hz) maintained a stable relationship with gene expression across all developmental stages, suggesting a consistent role in sustaining functional integrity throughout life.

These findings highlight the dynamic, frequency-specific nature of the molecular-genetic mechanisms underlying neurophysiological individuality. While the overarching genetic basis remains consistent, the specific contributions of different frequency bands evolve to meet changing functional demands and developmental priorities.

## Discussion

A growing body of research demonstrates that brain activity patterns, much like a fingerprint, are unique to each individual^9–12^. These neurophysiological traits are heritable^19^, correlate with inter-individual differences in cognition^10,11,14^, and manifest in disease processes^15,16,59^. However, the developmental trajectory of these traits over the lifespan has been relatively unexplored. This study addressed a fundamental question: Do individual neurophysiological traits become more distinctive across key developmental and aging stages?

Our findings reveal that individuals can be accurately differentiated based on patterns of periodic brain activity across the lifespan. However, the cortical regions most responsible for this differentiation shift with age. While neurophysiological traits are more homogeneous in childhood, sensorimotor regions become increasingly distinctive during the transition to adulthood. This shift is accompanied by a growing alignment between neurophysiological activity and gene expression, particularly involving genes related to ion transport and neurotransmission.

### Periodic Brain Activity as a Hallmark of Individuality

This study underscores the centrality of periodic brain activity in distinguishing individuals across all age groups (Figure S9). This finding is consistent with prior research showing that periodic features remain robust markers of individuality, even in conditions such as Parkinson’s disease, where aperiodic features lose their self-similarity^16^.

Notably, children (ages 4-11) exhibited differentiation accuracy nearly equivalent to that of adolescents and adults, despite the rapid and dynamic developmental changes occurring in childhood. This suggests that stable, individual-specific neurophysiological traits emerge early in life, even amid ongoing structural and functional maturation. The distinctiveness of children’s periodic traits aligns with the development of major brain rhythms, including the transition from lower frequencies (3-7 Hz) to the alpha rhythm (8-13 Hz)^28–31^, which stabilizes in late childhood, with a major developmental turning point around age seven^68^. Similar patterns are observed in somatomotor rhythms^69^, suggesting that neurophysiological individuality is present even as these rhythms continue to mature. These results challenge the assumption that younger brains are less individualized and highlight the resilience of periodic activity as a foundation for neurophysiological models.

Interestingly, while differentiation accuracy remained high across all age groups, other-similarity (similarity between an individual’s traits and those of others) increased with age for periodic traits (Figure S9). Conversely, aperiodic traits became increasingly distinct with age (Figure S8). These opposing patterns suggest that periodic traits provide a stable marker of individuality, while aperiodic traits may evolve adaptively, possibly reflecting cortical plasticity and functional integration over the lifespan. Future research should explore the interplay between these components across cognitive and behavioral contexts.

### Gene Expression and the Molecular Basis of Neurophysiological Differentiation

Our results shed light on how the alignment between cortical gene expression and individualized neurophysiological traits evolves across the lifespan (Figure 3). This alignment progressively strengthens with age, peaking in late adolescence, suggesting an increasing role of genetic factors in shaping brain individuality during neurodevelopment. Genes related to ion transport and neurotransmission appear to drive this alignment, emphasizing the genetic underpinnings of individualized brain activity.

We observed that specific frequency bands contribute differentially to gene-neurophysiology alignment. Alpha band activity (8-13 Hz) consistently aligned with positive gene signatures across all ages, suggesting its enduring role in maintaining functional integrity. Conversely, theta band activity (4–8 Hz) showed an association with neurogenesis-related genes during early development, which diminished later in life. Beta band activity (13–30 Hz), associated with predictive coding and motor functions, demonstrated increasing alignment with positive gene signatures into adulthood. These findings illustrate the dynamic nature of frequency-specific molecular mechanisms underlying neurophysiological individuality. They may mirror the brain’s changing functional demands across the lifespan. For instance, the increasing alignment of beta activity with genes involved in ion transport and neurotransmission in adulthood may reflect the greater reliance on sensorimotor networks for adaptive behaviour.

Further, the distinct roles of positive and negative gene signatures highlight the complexity of brain individuality. While the positive signature consistently aligned with differentiable neurophysiological traits, the negative signature—linked to emotional processing^19,70^—showed greater variability across age groups. This suggests that different molecular pathways govern neurophysiological stability, and flexibility, with positive genes supporting cognitive and motor functions, and negative genes driving emotional regulation and responses to environmental stress.

### The Increasing Role of Sensorimotor Regions in Differentiation Across the Lifespan

Our findings show that the cortical regions most critical for individual differentiation shift over time, with sensorimotor regions playing an increasingly prominent role as individuals mature into adulthood (Figure 2a). This shift aligns with research indicating that sensorimotor areas gain importance in functional brain organization as motor control and sensorimotor integration mature during neurodevelopment^64^.

Additionally, the alignment between cortical regions and functional gradients evolves with age. In childhood, characteristic brain regions align more closely with the visual-to-motor gradient, which plays a key role in early brain development. In adulthood, this alignment shifts toward the unimodal-to-transmodal gradient, reflecting the maturation of higher-order association cortices responsible for complex cognitive and emotional functions^24,25,46,71–74^ (Figure 2b).

This developmental shift underscores the role of sensorimotor regions in supporting predictive coding and inferential models, processes fundamental to cognition^48–50,75–78^. The increasing prominence of beta activity in these regions further highlights their genetic and functional specialization during development^19^.

### Neurophysiological Differentiation in Older Adults: A Compensatory Mechanism?

Our findings also reveal greater differentiation in older adults based on broadband neurophysiological spectral features originating from sensorimotor regions, including motor and ventral-medial visual areas. This may reflect compensatory mechanisms for age-related sensory decline, as older adults often exhibit reduced neural responsiveness to sensory stimuli^79–81^. Such compensatory strategies likely vary across individuals, potentially explaining the increased distinctiveness of sensorimotor activity in older adults. Future studies should investigate the mechanisms underlying these processes and their contribution to neurophysiological differentiation in aging populations.

### Implications for Health Monitoring and Brain-Behaviour Relationships

Our findings underscore the importance of developmental stages in understanding neurophysiological differentiation and highlight the need for multi-omic data incorporating diverse socioeconomic, age, and geographical backgrounds^82^. Extending the concept of paediatric growth charts^20^ to neurophysiological traits could enable novel approaches to monitoring brain health for tracking age-related brain changes. Extending this concept to neurophysiological traits could create novel opportunities for monitoring brain health and detecting deviations indicative of neurological or psychiatric conditions^84^.

The stability of both periodic and aperiodic traits has implications for understanding neurodevelopmental and neurodegenerative disorders. Delayed stabilization of these traits has been linked to atypical neurodevelopment, while greater stabilization correlates—albeit weakly— with better cognitive outcomes^54^. By establishing age-matched normative variants for neurophysiological differentiation, our study provides a foundation for further research into the diagnostic and prognostic utility of these traits in clinical contexts.

### Limitations and Future Directions

While our study included a large, cross-sectional lifespan sample, longitudinal data are needed to provide a more accurate depiction of how neurophysiological traits evolve within individuals over time. Additionally, our gene expression analyses relied on an adult dataset, limiting insights into gene-neurophysiology alignment across developmental stages. Future research should prioritize data collection in infancy and early childhood, which remain underexplored. Advances in optically pumped magnetometers (OPMs) may facilitate neurophysiological data acquisition in younger populations^83^.

## Conclusion

Our findings demonstrate that neurophysiological traits are unique to individuals but evolve throughout the lifespan. These traits reflect dynamic interactions between genetic and environmental influences, with sensorimotor regions playing an increasingly prominent role in differentiation. By considering developmental and aging trajectories, future research can better capture the dynamic nature of the neurophysiological self and its implications for understanding individuality and brain health.

## Methods

### Participants: SickKids Cohort

Data were collected from 401 healthy individuals aged 4-68 years (mean age = 20.79, SD = 14.14; 169 female). Resting-state eyes-open magnetoencephalography (MEG) recordings lasted approximately 5 minutes and were acquired using a 151-channel whole-head CTF MEG system (Port Coquitlam, British Columbia, Canada) sampled at 600 Hz. Participants’ head positions were continuously monitored throughout the recording using three Head-Position Indicator (HPI) coils to ensure consistent head localization. All participants also underwent structural T1-weighted MRI to inform MEG source modelling.

### Participants: Cam-CAN Cohort

Data were collected from 606 healthy individuals aged 18-89 years (mean age = 54.69, SD = 18.28; 299 female) from the Cambridge Centre for Aging and Neuroscience repository (CamCAN)^62^. Each participant underwent a resting-state, eye-closed MEG recording using a 306-channel VectorView MEG system (MEGIN, Helsinki, Finland). The MEG system consisted of 102 magnetometers and 204 planar gradiometers, sampled sampling rate of 1 kHz with a 0.03-330 Hz bandpass filter. Resting-state recordings lasted approximately 8 minutes. Continuous monitoring of participants’ head positions was performed using four HPI coils. Additionally, electrooculography (EOG) and electrocardiography (ECG) electrodes were used to capture ocular and cardiac artifacts for subsequent removal. T1-weighted MRI scans were also collected from all participants.

For demographic details of each dataset, see Supplemental Tables S1 and S2.

### Preprocessing of MEG Data

MEG data preprocessing was conducted using Brainstorm^84^ (version dated 08-08-2023) in MATLAB R2021a (Mathworks Inc., Natick, Massachusetts, USA) adhering to good practice guidelines^85^. Preprocessing followed our prior work on MEG individual differentiation^9,16,19^.

Line noise artifacts at 50 Hz (CamCAN) and 60 Hz (SickKids), along with their first ten harmonics, were removed using a notch filter bank to ensure the removal of environmental and power-line interference. Additionally, an 88 Hz artifact characteristic of the CamCAN dataset was removed.

Slow-wave artifacts and DC offsets were mitigated using a high-pass finite impulse response filter with a 0.3-Hz cut-off frequency in both datasets.

### Source Modeling of MEG Data

Brain source models were derived using individual participants’ T1-weighted MRI scans to constrain MEG source mapping. MRI volumes were segmented and labelled automatically using FreeSurfer (version 7.3.2)^86^ . The MRI data were co-registered with MEG recordings using approximately 100 digitized head points, when available.

We used Brainstorm’s overlapping-spheres approach with default parameters for individual head models. Cortical source models were then computed with linearly-constrained minimum-variance (LCMV) beamforming (Brainstorm 2018 version for source estimation processes). MEG source orientations were constrained to be normal to the cortical surface. A grid of approximately 15,000 locations across the cortex was used for source modelling.

### Neurophysiological Traits

The power spectrum of MEG source time series at each cortical location was computed using the Welch method, with 2-second time windows overlapping by 50%. Cortical surfaces were parcellated into 148 cortical regions using the Destrieux atlas^87^. We excluded the delta band (1–4 Hz) in the SickKids dataset due to the low signal-to-noise ratio, limiting analysis to frequencies between 4–150 Hz. For CamCAN, the analysis covered 1–150 Hz. The frequency resolution was set to 0.5 Hz, resulting in matrices of 148 cortical regions by 293 (SickKids) or 300 (CAmCAN) frequency features for each participant. Spectral features were then exported to the R (version 4.2.2)^88^ for individual differentiation analyses.

### Differentiability

Our differentiation method was adapted from previous work on neurophysiological differentiation^9–11^. Neurophysiological differentiation relied on differentiability of each participant across resting-state segments. We divided recordings into first and second halves to evaluate reproducibility and distinctiveness.

We obtained Pearson correlations between participant_i_’s traits from the first segment and all traits at the second segment. Correct differentiation occurred if participant_i_’s self-similarity was greater than their other-similarity with other participants. Differentiation accuracy was defined as the percentage of correctly differentiated participants across the cohort.

We defined differentiation accuracy as the percentage of correctly differentiated participants across the entire cohort. For the SickKids cohort, we computed the differentiation accuracy in children (N = 148; 4-11 years old), adolescents (N = 57; 11-18 years old), adults (N = 196; 18+ years old), and the entire cohort (N = 401). We chose these age group boundaries to maximize the number of participants in the children and adult groups. To verify the robustness of our results, we slightly adjusted the definitions of the age groups and assessed their impact on differentiation accuracy. Details of this robustness check are presented in Figure S7.

A similar approach was used for the CamCAN cohort: young adults (18-45 years, N = 204), middle-aged adults (45-65 years, N = 194), older adults (65+ years, N = 208), and the entire cohort (N = 606).

Additionally, we computed a continuous differentiability score for each participant^9^. This score was calculated by z-scoring self-similarity relative to the mean and standard deviation of other-similarity scores. A high differentiability score indicated that a participant’s traits were more distinct from others in the cohort.

### Bootstrapping Differentiation Accuracy

To assess reliability, we derived bootstrapped 95% confidence intervals for each differentiation accuracy. Participants were randomly sampled with replacement, subsampling 485 participants for the entire cohort and 175 for each age group in the CamCan sample. For the SickKids Institute sample, 361 participants for the entire cohort were subsampled. Bootstrapping was repeated 1,000 times, and 2.5^th^ and 97.5^th^ percentiles of the resulting accuracy distribution were computed to provide confidence intervals, which reflect the empirical uncertainty.

The specific choice of the number of subsampled participants was made to balance the computational load with statistical power while ensuring consistency across the bootstrap samples. By defining our approach in this way, we aimed to maintain an adequate sample size that ensures robust estimation of the true differentiation accuracy for each cohort while accommodating practical considerations for data processing.

### Band-limited Neurophysiological Traits

We replicated differentiation analyses using canonical electrophysiological frequency bands: delta (1–4 Hz), theta (4–8 Hz), alpha (8–13 Hz), beta (13–30 Hz), gamma (30–50 Hz), and high gamma (50–150 Hz). Delta band activity was excluded from SickKids analyses due to poor signal-to-noise ratio (SNR), limiting reliable differentiation.

Each canonical frequency band was subjected to the same differentiation analysis.

### Recording Artifacts and Differentiability

We assessed the effect of common physiological artifacts on differentiability using regression models. Artifact levels were quantified using root-mean-square (RMS) values of ocular (VEOG, HEOG), cardiac (ECG), and head movement (HLU channels) artifacts across the MEG recording. Analyses were conducted separately for each dataset to account for specific differences.

We used regression models to determine whether these physiological artifacts influenced individual differentiability. These analyses were conducted separately for the SickKids and CamCAN datasets to account for cohort-specific variations in recording environments, MEG system used, participant characteristics, and artifact prevalence.

In the SickKids cohort, we further assessed the impact of total intracranial volume (TIV), as head size may vary significantly among children. TIV was calculated using FreeSurfer, and regression analyses were used to determine if it influenced differentiability.

### Empty-Room Differentiation

To establish that individual differentiation was not driven by environmental or instrumental noise, we conducted empty-room differentiation using MEG recordings from the day of each participant’s visit. For the SickKids dataset, only a single empty-room recording was available for the entire cohort. This recording was used to approximate the baseline noise level. Empty-room data were pre-processed similarly to participant data, except for physiological artifact removal. These recordings were used to generate pseudo neurophysiological traits. Differentiation accuracy using these pseudo-traits served as a baseline performance measure, ensuring that results were not due to noise^9^.

### Aperiodic vs. Periodic Spectral Parametrization

To assess the contribution of aperiodic neurophysiological activity to individual differentiability, we parametrized the participants’ MEG source power spectra using the ms-specparam tool in Brainstorm^84,89^. We extracted the aperiodic background components from periodic oscillatory peaks in each source signal spectrum.

For the SickKids dataset, the frequency range for analysis was set between 4 and 50 Hz, while for the Cam-CAN dataset, it was set between 1 and 40 Hz. The peak width limits differed slightly, with the SickKids dataset having limits between 1 and 8 Hz, and the Cam-CAN dataset between 0.5 and 12 Hz. We specified a maximum of six peaks per spectrum, with a minimum peak amplitude of 1 arbitrary unit (a.u.). The peak detection threshold was set at two standard deviations above the mean, while the proximity threshold, determining how close detected peaks could be, was set at 0.75 standard deviations. The aperiodic component was modelled using a fixed mode to ensure consistency across participants.

These hyperparameters for spectral parametrization were determined based on visual inspection of the spectra, to ensure that the model captured the relevant spectral features while minimizing the risk of overfitting or misinterpretation.

Following spectral parametrization, the (a)periodic components were used to derive neurophysiological traits defined from (a)periodic spectral features. The differentiation procedure was then carried out using the previously described approach, where we evaluated how well participants could be differentiated based solely on the aperiodic vs. periodic spectral components fits of their cortical neurophysiological activity.

### Relative Contribution of Neurophysiological Traits

We calculated intraclass correlations (ICC) to evaluate the contribution of each feature (frequency x cortical region) to participant differentiation^9,10,90^. High ICC values indicate that a specific neurophysiological feature is consistently similar within participants across repeated measures while being distinct across different participants. Therefore, features with high ICC values contribute most significantly to participant differentiation.

To generate the brain maps presented in Figures 2 and S11, we initially averaged ICC values within each canonical frequency band (e.g., theta, alpha, beta, gamma). Following this, we averaged across the resulting frequency band maps to derive a broadband saliency topography. This two-step averaging was performed to ensure that each frequency band had an equal influence on the overall saliency map, independent of differences in frequency band definitions (e.g., the theta bandwidth spans 4 Hz, while gamma covers at least 20 Hz).

We pooled data from both the SickKids and CamCAN datasets to obtain a comprehensive representation of how feature saliency changes across the lifespan. Participants were ordered by ascending age, and we used a sliding window approach to calculate ICC values for each age group. Specifically, we selected a moving window of 100 participants with 75% overlap between consecutive windows, which resulted in 37 ICC maps representing increasing age bins. This method allowed us to capture the gradual changes in the contribution of different cortical regions and frequency bands to individual differentiation as participants age. The results of this analysis are visualized in Figure 2a, which illustrates the shifting topographic features that characterize individualized neurophysiology across the lifespan.

### Neuroanatomical Similarity

To ensure that increased other-similarity observed in children was not due to anatomical features, we conducted additional analyses incorporating neuroanatomical data. Specifically, we extracted nine anatomical features for each cortical region defined by the Destrieux atlas^87^, which were provided by the FreeSurfer segmentation process^86^. These features included metrics such as cortical thickness, surface area, and volume.

We used these anatomical features to construct an anatomical similarity matrix that quantified the similarity between each pair of participants based on their anatomical characteristics. This matrix was derived by computing Pearson correlations between the anatomical features of all possible participant pairs.

We then evaluated whether this anatomical similarity could explain the observed other-similarity in neurophysiological traits, particularly the greater similarity observed in children. To do this, we computed the linear relationship between participant-pair anatomical similarity and neurophysiological similarity across all pairs.

### Neurophysiological Traits and Cortical Functional Hierarchy

We assessed whether the pattern of cortical regions contributing most to differentiation increasingly aligned with the functional organization of the cortex with age. Specifically, we evaluated their correspondence with the unimodal-to-transmodal gradient, which represents the functional hierarchy of cortical regions from primary sensory to higher-order association areas^63^.

To evaluate this correspondence, we computed the spatial alignment between the ICC maps (derived from the sliding window approach, see above) and the unimodal-to-transmodal functional gradient obtained using the neuromaps toolbox^91^. This alignment was quantified using Pearson correlation for each age bin, capturing the degree to which regions involved in individual differentiation align with the functional gradient of the cortex.

We further examined whether this alignment changed non-linearly across the lifespan by fitting a third-order polynomial model to capture potential trends. To test the robustness of observed changes, we used 1,000 autocorrelation-preserving permutation tests using the Hungarian spin method. These permutation tests generated spin-based resampling of cortical maps to assess statistical significance while preserving spatial autocorrelation^92,93^.

For each spin permutation, we recalculated the alignment between the permuted functional gradient map and the ICC maps, fitting a third-order polynomial to the resulting alignment data. We then computed the permuted p-value by comparing the observed third-order polynomial beta-coefficients with those obtained from the permuted data, allowing us to assess the statistical significance of the observed alignment patterns.

We also evaluated the alignment between the pattern of cortical regions contributing the most to differentiation and the visual-to-motor functional gradient^63^, hypothesized to be more relevant for early brain development. The same procedures for spatial alignment and permutation testing were applied (see Figure 2a).

### Gene Expression

We retrieved cortical gene expression data from six postmortem brains from the Allen Human Brain Atlas (AHBA; http://human.brain-map.org/)^67^. Data were processed using the abagen Python package^94^, following our previously established pipeline^19^, with the exception of using the Destrieux atlas for cortical parcellation. We selected microarray probes with the highest differential stability to represent the expression of each gene, resulting in 20,232 genes included in our analysis.

Tissue samples were assigned to the nearest cortical region using a nonlinear registration method, focusing on minimizing misalignment across regions. Gene expression data were normalized across tissue samples and subjects^95^, and only genes with differential stability greater than 0.1 were retained, resulting in a final set of 9,278 genes^70,94,96,97^.

We used previously defined sets of positive and negative genes, categorized based on their association with participant differentiation^19^. Genes positively correlated with differentiation were considered positive, while those negatively correlated were negative. After filtering for stability, we retained 2,076 positive genes and 2,219 negative genes. Cortical maps representing gene expression were generated to examine their alignment with the differentiable neurophysiological features observed across the lifespan.

### Topographic Similarity of Gene Expression Across the Lifespan

To evaluate the stability of positive and negative gene signatures identified in our previous work^19^, we analyzed gene expression data across developmental stages, from 8 post-conception weeks to 40 years of age, using data from 16 cortical regions obtained from the BrainSpan Atlas^98^. Gene expression data were grouped into five life stages^19,70,99^: fetal (8–37 post-conception weeks), infant (4 months–1 year), child (2–8 years), adolescent (11–19 years), and adult (21–40 years). For each developmental stage, we computed the average expression levels of the positive and negative gene sets across the cortical regions.

To assess the stability of these spatial gene expression patterns across different life stages, we calculated Pearson correlations between the cortical expression patterns of all pairs of life stages. This resulted in a symmetric topographic similarity matrix for the positive and negative gene sets separately (Figure S14).

### Correspondence of Neurophysiological Traits with Cortical Gene Expression

To investigate whether the alignment between the pattern of cortical regions that contribute the to differentiation and cortical gene expression changed across the lifespan, we computed the spatial alignment between the ICC maps and the positive gene expression signature. This alignment was assessed using Pearson correlations to quantify the similarity between the spatial distribution of differentiable features and gene expression patterns.

We used a sliding window approach across different age bins to examine changes in this alignment throughout the lifespan. This allowed us to explore how the developmental progression of neurophysiological features aligns with the expression of genetic systems across the cortex.

### Partial Least Squares Analysis

To further explore the alignment between cortical gene expression and neurophysiological traits, we conducted a Partial Least Squares (PLS) analysis for each age group. This multivariate analysis was used to relate the most differentiable cortical regions (as quantified by ICC values) at each age bin (Figure 2a) to cortical gene expression patterns obtained from the Allen Human Brain Atlas^67^.

For each age bin, we constructed two data matrices: one containing distinctive neurophysiological traits (represented by the ICC values) and another containing the cortical gene expression data. The traits matrix had four columns (each representing a frequency band) and 148 rows, corresponding to the cortical regions of the Destrieux atlas. The gene expression matrix had 9,278 columns (representing genes) and 148 rows (representing cortical regions).

To ensure comparability, we z-scored the columns of each matrix before applying Singular Value Decomposition (SVD) to the cross-covariance matrix between the neurophysiological and gene matrices. The resulting latent components provided insight into the covariance structure between the neurophysiological features and gene expression across age bins:

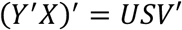

Here, *U* is a 9,278 by 4 orthonormal matrix, and *V* is a 4 by 4 orthonormal matrix, with each column representing a latent component of the covariance between neurophysiological traits and gene expression. This decomposition was repeated for each age group using a sliding window approach, as described above. We reported the percentage of covariance explained by each latent component, giving insight into the relationships between neurophysiological features and genetic expression across age bins^100,101^.

To assess the significance of the latent components, we conducted 1,000 spatial autocorrelation-preserving permutation tests using the Hungarian spin method (detailed above). We generated null distributions of singular values from these permutations to compute empirical p-values, which were corrected for multiple comparisons using False Discovery Rate (FDR) correction, as implemented in MATLAB.

To determine the contribution of individual genes and frequency bands to the observed patterns of covariance, we computed Pearson correlations between each variable’s spatial distribution over the cortex (i.e., gene expression and ICC values) and the opposing PLS brain score pattern^19,70^. These loadings are bounded between -1 and 1, facilitating intuitive interpretation—large absolute loadings indicate strong contributions to the latent component of covariance.

Lastly, to evaluate the consistency of gene contributions across the lifespan, we assessed the similarity between the gene loadings derived from each age bin with our previously reported gene loadings from an independent PLS analysis using MEG data from the Human Connectome Project^19^. This was done using Pearson correlations (Figure 3b, right panel), which provided insight into the stability of the genetic signature governing differentiable neurophysiological activity throughout development and aging.

## Data availability

Data used in the preparation of this work are available through the CamCAN repository (https://camcan-archive.mrc-cbu.cam.ac.uk/)^102^. Data from the SickKids cohort will be available upon reasonable request. Cortical gene expression data is available from the AHBA at https://human.brain-map.org/static/download, and the BrainSpan database at https://www.brainspan.org/static/download.html.

## Code availability

All codes for preprocessing, data analysis, and data visualization can be found on the project’s GitHub.

## Acknowledgements

Funding to SB included a Discovery grant from the Natural Science and Engineering Research Council of Canada (436355-13), the CIHR Canada Research Chair in Neural Dynamics of Brain Systems (CRC-2017-00311), and from the NIH (R01-EB026299-05). This work was also supported by a doctoral fellowship from NSERC (JDSC) and to AIW by a CIHR Banting Postdoctoral Fellowship (BPF-186555) and the Canada Research Chair (CRC-2023-00300) in Neurophysiology of Aging and Neurodegeneration. The SickKids MEG data collection was supported by CIHR funding to MJT (MOP 106582, 137115, 142379, 119541).

## Supplemental Material

### Datasets and Demographics

The study relied on MEG recordings from two datasets, as detailed in the main text: the SickKids and CamCAN datasets^62^. Below, we provide a comprehensive summary of relevant demographic variables for both datasets.

**Table S1.**
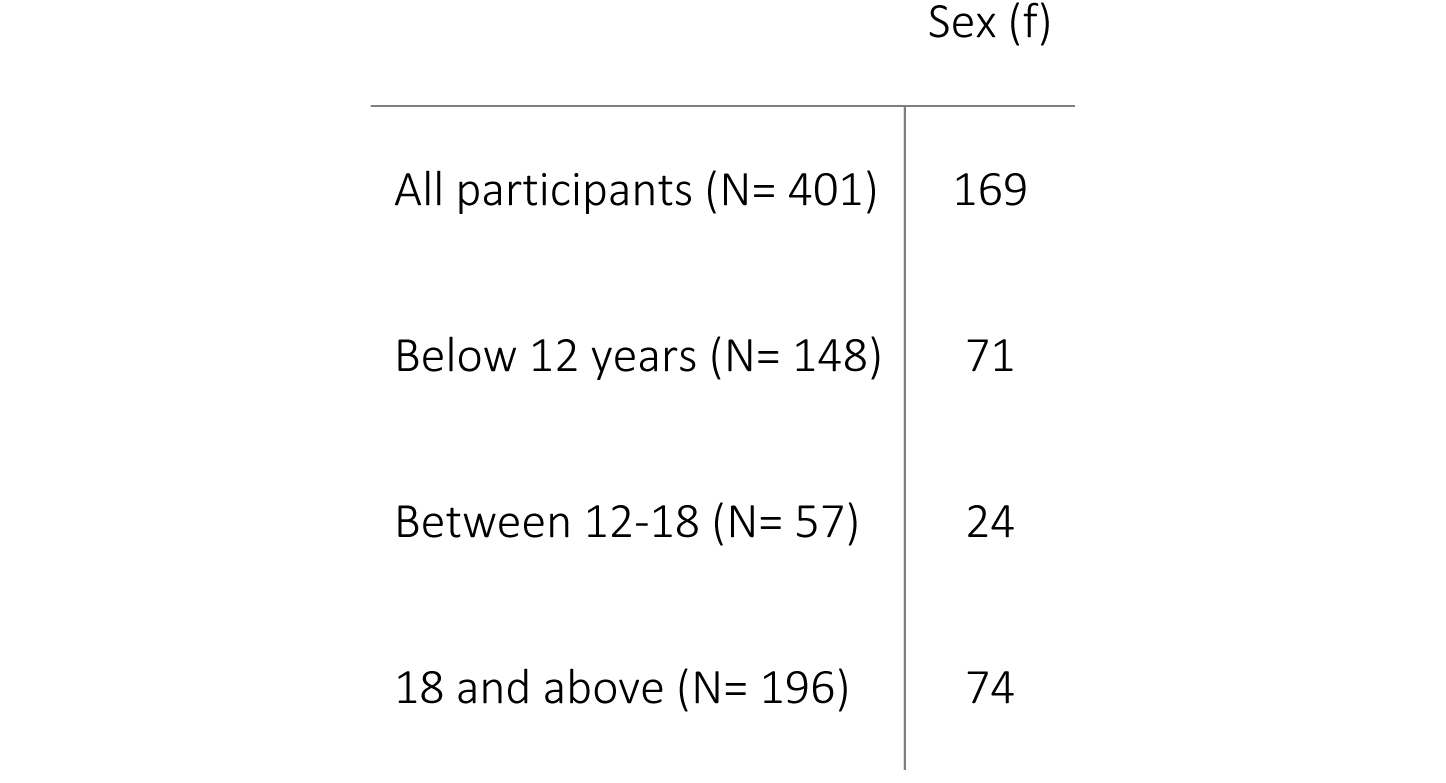
Demographics of SickKids participants.

**Table S2.** Demographics of CamCAN participants.

|  | Sex (f) | Handedness (sd) | MMSE (sd) |
| --- | --- | --- | --- |
| All participants N=606 | 299 | 78.9 (49.2) | 28.95 (1.25) |
| Young adults (18-45) N=204 | 103 | 75.3 (52.5) | 29.3 (1.05) |
| Adults (45-65) N=194 | 100 | 76.6 (53.3) | 29.1 (1.07) |
| Older adults (65+) N=208 | 96 | 84.9 (40.8) | 28.4 (1.38) |

**Figure S1:**
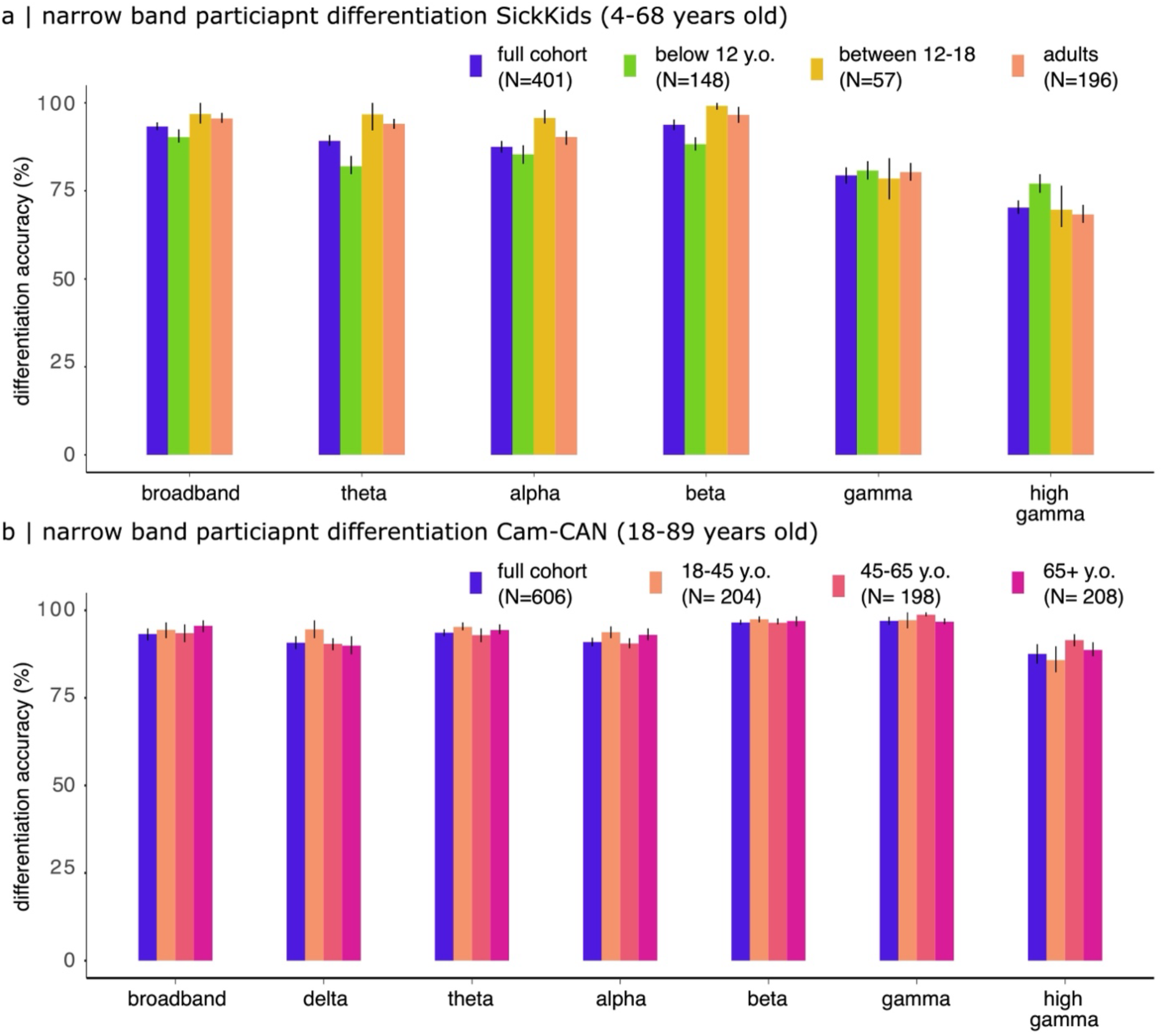
Frequency Band-Specific Differentiation. a) Participant differentiation accuracy in the SickKids cohort: Accuracy in distinguishing participants based on their neurophysiological traits derived from narrow-band frequency components, estimated using 2-minute MEG data segments. Error bars represent bootstrapped 95% confidence intervals. (b) Participant differentiation accuracy in the CamCAN cohort: Accuracy of participant differentiation using narrow-band neurophysiological traits from the CamCAN dataset. Narrow-band analyses reveal the contributions of individual frequency bands to differentiation across participants. Error bars denote bootstrapped 95% confidence intervals.

**Figure S2.**
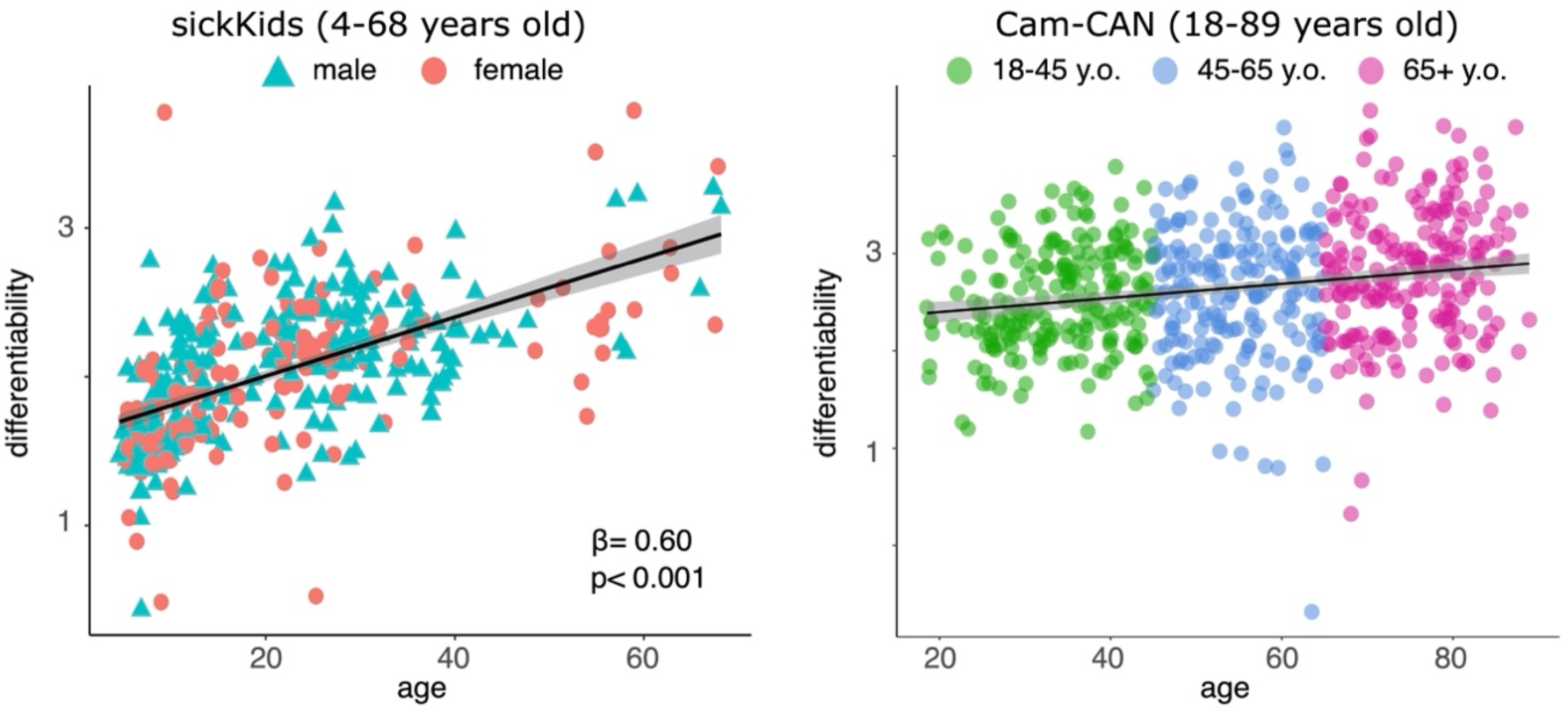
Decreased Differentiability of Children from Wideband Neurophysiological Traits. Younger individuals are less easily differentiated based on their neurophysiological traits compared to older individuals.

- Left panel (SickKids dataset): A strong linear relationship is observed between age and differentiability, explaining 36% of the variance. Colours represent biological sex in this cohort.
- Right panel (CamCAN dataset): A weaker linear relationship is evident in adults, explaining only 4% of the variance. Colours indicate age groups within the CamCAN dataset. These findings suggest that the ease of differentiation increases with age, particularly during childhood and adolescence.

### Sensitivity Analyses Against Artifacts (CamCAN)

We conducted sensitivity analyses to ensure that our results were not influenced by environmental or physiological artifacts.

#### 1. Environmental Factors

To evaluate the impact of environmental noise on participant differentiation, we used empty-room MEG recordings to define pseudo neurophysiological traits. These recordings were processed identically to the participants’ data and projected onto their cortical surfaces using the same imaging pipeline. This resulted in a set of pseudo neurophysiological traits defined solely by noise. Differentiation accuracy based on these pseudo-traits was minimal (<5%), confirming that environmental noise did not significantly contribute to participant differentiation (see grey bars in Figure 1b).

#### 2. Physiological Artifacts

To assess the influence of physiological artifacts, we analyzed relationships between participant differentiation and head motion, cardiac, and ocular artifacts:

`o` **Head Motion**: Older adults exhibited greater head motion, with a significant linear increase observed with age (t(604) = 9.22, r = 0.35, p < 0.001; Figure S3).
`o` **Cardiac Artifact**: A significant linear decrease in cardiac artifact was noted with age (t(604) = -4.94, r = -0.20, p < 0.001; Figure S3).
`o` **Ocular Artifact**: No significant relationship between age and ocular artifacts was detected (t(604) = 0.27, r = 0.01, p = 0.78; Figure S3).

Crucially, none of these artifacts were significantly related to differentiability:

`o` **Head Motion**: r = 0.06, p = 0.13
`o` **Ocular Artifacts**: r = 0.00, p = 0.92
`o` **Cardiac Artifact**: r = -0.03, p = 0.45

These findings suggest that while older adults exhibited greater head motion artifacts, this did not affect participant differentiation. Similarly, neither cardiac nor ocular artifacts impacted differentiability, further validating the robustness of our results against physiological noise.

**Figure S3.**
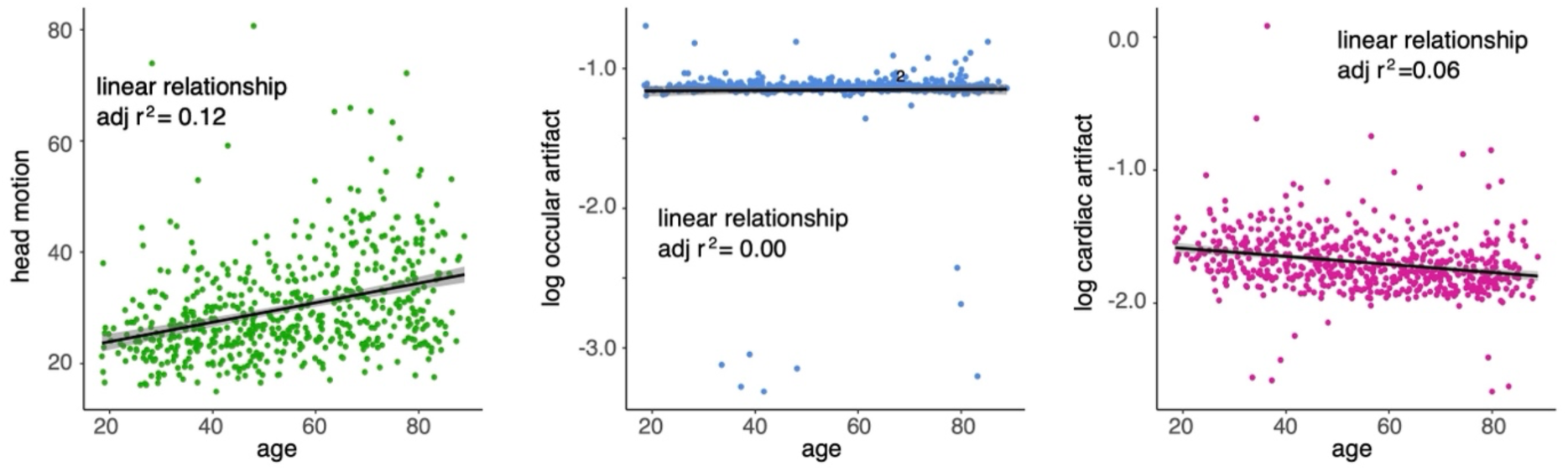
Influence of Physiological Artifacts (Cam-CAN Dataset). We examined the linear relationships between chronological age and common physiological artifacts in the CamCAN dataset:

- Left panel: A moderate positive linear relationship was observed between head motion and age, indicating that older participants exhibited increased head motion during MEG recordings.
- Middle panel: Ocular artifacts showed no meaningful relationship with age, explaining 0% of the variance.
- Right panel: Cardiac artifacts exhibited a minimal relationship with age, explaining 6% of the variance. These findings suggest that while head motion increases with age, other physiological artifacts like ocular and cardiac signals contribute minimally to age-related variance in MEG recordings.

### Sensitivity Analyses Against Artifacts (SickKids)

We assessed the influence of cranial volume and head motion—two common physiological confounds in MEG recordings—on participant differentiation in the SickKids cohort.

1. **Cranial Volume**: As expected, cranial volume exhibited a significant non-linear increase with age (β = -1.61 × 10⁵, SE = 2.37 × 10⁴, 95% CI [-2.07 × 10⁵, -1.14 × 10⁵], p < 0.001; Table S3 and Figure S4). This trend reflects normal developmental changes during childhood and adolescence.
2. **Head Motion**: Head motion decreased significantly with age in a non-linear manner (β = 3.39, SE = 0.39, 95% CI [2.63, 4.15], p < 0.001; Table S4 and Figure S4), consistent with improved motor control and behavioral regulation in older participants.
3. **Impact on Differentiability**: Despite these age-related changes in cranial volume and head motion, neither significantly influenced the relationship between age and differentiability. When cranial volume and head motion were included as nuisance covariates in a linear regression model, the relationship between age and differentiability remained robust (β = 0.02, SE = 1.43 × 10⁻³, 95% CI [0.01, 0.02], p < 0.001; Table S5 and Figure S4).

These findings suggest that while cranial volume and head motion vary significantly with age, their impact on participant differentiation is minimal. Differentiability remains largely driven by age-related neurophysiological changes rather than physiological artifacts, further supporting the reliability of our results.

**Figure S4.**
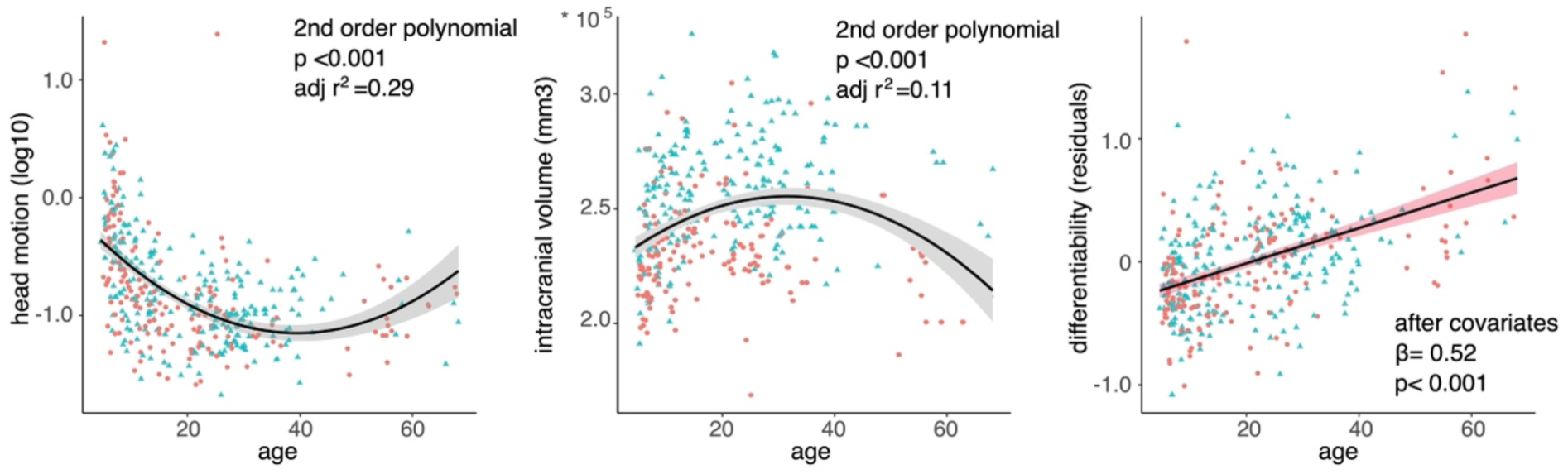
Confounds and Artifacts (SickKids Dataset). We analyzed the relationships between chronological age and physiological artifacts in the SickKids dataset:

- Left panel: Head motion exhibited a non-linear decrease with age, reflecting reduced movement during MEG recordings as participants aged.
- Middle panel: Intracranial volume increased non-linearly with age, consistent with expected patterns of cranial growth in childhood and adolescence.
- Right panel: Physiological artifacts (e.g., head motion and intracranial volume) had minimal impact on the linear relationship between age and differentiability, confirming the robustness of age-related differentiation patterns to these potential confounds.

**Table S3.** Intracranial Volume Increases With Age Nonlinearly (SickKids).

| Intracranial volume |  |  |  |
| --- | --- | --- | --- |
| Predictors | Estimates | CI | p |
| (Intercept) | 245766.70 | 243429.93 – 248103.47 | <0.001 |
| Age [1st degree] | 39341.25 | -7414.60 – 86097.10 | 0.099 |
| Age[2nd degree] | -161096.32 | -207805.63 – -114387.01 | <0.001 |
| Observations | 399 |  |  |
| R<sup>2</sup> / R<sup>2</sup> adjusted | 0.109 / 0.105 |  |  |

**Table S4.** Head Motion Decreases with Age Nonlinearly (SickKids).

| Log<sub>10</sub> head motion |  |  |  |
| --- | --- | --- | --- |
| Predictors | Estimates | CI | p |
| (Intercept) | -0.79 | -0.83 – -0.76 | <0.001 |
| Age [1st degree] | -3.74 | -4.51 – -2.98 | <0.001 |
| Age [2nd degree] | 3.39 | 2.63 – 4.15 | <0.001 |
| Observations | 401 |  |  |
| R<sup>2</sup> / R<sup>2</sup> adjusted | 0.298 / 0.295 |  |  |

**Table S5.** Increased Differentiability with Age is Robust Against Physiological Artifacts (SickKids).

| Differentiability |  |  |  |
| --- | --- | --- | --- |
| Predictors | Estimates | CI | p |
| (Intercept) | 0.00 | -0.08 – 0.08 | 0.954 |
| Age | 0.52 | 0.43 – 0.60 | <0.001 |
| volume | 0.05 | -0.03 – 0.13 | 0.185 |
| log10(head motion) | -0.18 | -0.27 – -<br>0.10 | <0.001 |
| Observations | 399 |  |  |
| R<sup>2</sup> / R<sup>2</sup> adjusted | 0.387 / 0.382 |  |  |

To further evaluate the robustness of our results, we defined a subset of children (under 12 years old; n = 57) matched to adults in terms of both intracranial volume and head motion artifact.

1. **Matching Process**

a. **Intracranial Volume**: The children in this subset were matched to adults based on intracranial volume (permutation t-test: t = -2.02, p = 0.05).
b. **Head Motion**: No significant difference in head motion was observed between the two groups (t = 1.53, p = 0.12; Figure S5a).
2. **Differentiability Analysis**

a. Despite being matched on these confounding variables, children in the matched cohort were still more challenging to differentiate compared to adults. Differentiability remained significantly lower in children than in adults (permutation t-test: t = -3.76, p < 0.01; Figure S5b).
3. **Regional Topography**

a. The spatial distribution of the most characteristic brain regions for differentiating children remained highly consistent with the results reported in the main text (r = 0.77, p < 0.001, p_spin_ < 0.001; Figure S5c), demonstrating the reliability of our findings (see Figure 2a, far left, for comparison).

By controlling for intracranial volume and head motion artifacts, we confirmed that the observed disparities in differentiability between children and adults are not driven by these physiological factors. Additionally, the consistency in the topography of characteristic brain regions underscores the robustness of our differentiation approach across development.

**Figure S5.**
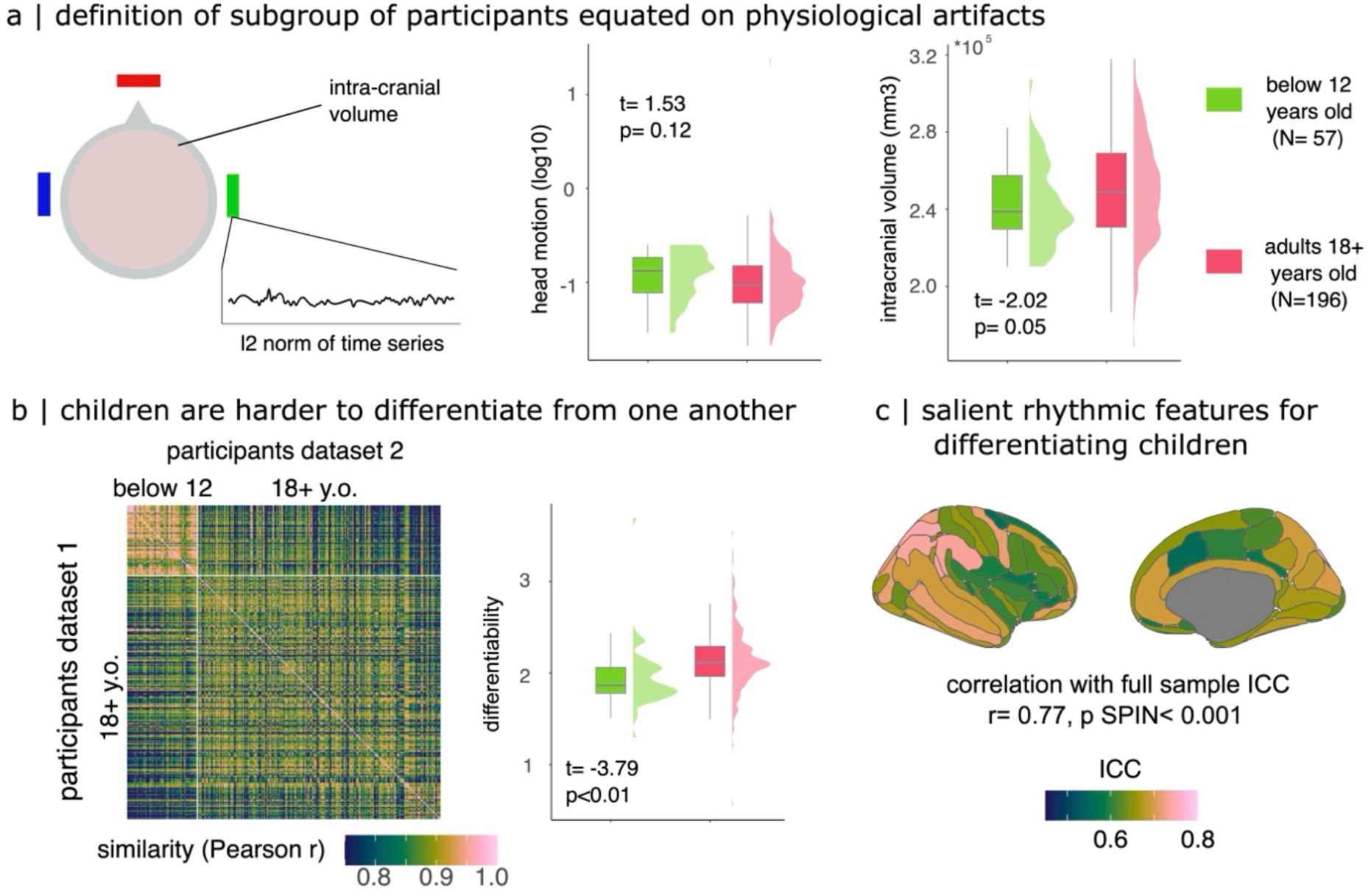
Weak Contribution of Physiological Artifacts to Differentiation. (a) **Physiological Artifact Matching**: A subgroup of children was defined to closely match adults in terms of head motion (middle panel) and intracranial volume (right panel). Box plots illustrate minimal differences in these physiological artifacts between the two groups, confirming comparability. (b) **Differentiability Challenges in Children**: **Left panel**: The similarity matrix, ordered by ascending age, highlights that children (indicated by the green square) are more challenging to differentiate due to higher neurophysiological traits similarity (off-diagonal elements). **Right panel**: A boxplot comparing participant differentiability shows that the children subgroup remains more difficult to differentiate compared to adults. (c) **Cortical Differentiation in Children**: Topographic maps of Intraclass Correlation Coefficients (ICCs) reveal the cortical regions most characteristic of individual differentiation in children. The brain map closely resembles the differentiation map for all children (Figure 2a), underscoring the robustness of these findings.

### Impact of Anatomy on Differentiation

To assess the influence of anatomical features on participant differentiation, we examined several cortical properties derived from FreeSurfer^86^ across parcels of the Destrieux atlas. These features included:

- Number of vertices,
- Grey matter volume,
- Curvature,
- Folding,
- Cortical thickness (mean and standard deviation), and
- Surface area.

#### Anatomical (Dis)similarity Matrix

We computed a participant dissimilarity matrix based on these anatomical features, analogous to the neurophysiological differentiation matrix depicted in Figure 1c. Interestingly, anatomical features showed different age-related patterns of similarity: The anatomical features of children were less similar to other children than the anatomical features of adults were to other adults (permutation t-test: t = -56.69, p < 0.01; Figure S6a).

#### Relationship Between Anatomy and Spectral Similarity

We explored the linear relationship between anatomical and spectral other-similarity across age groups. No meaningful linear relationships were observed:

- Children: r = -0.04
- Adolescents: r = 0.05
- Adults: r = -0.00

These findings were consistent when analyzing similarity derived from periodic neurophysiological traits (children: r = 0.06, adolescents: r = -0.06, adults: r = -0.01; Figure S6b).

#### Neurodevelopment and Anatomical Centile Scores

To explore how neurodevelopmental trajectories of brain structure impact participant differentiation, we leveraged anatomical brain-charts^20^ to calculate centile scores for each individual. These scores benchmark cortical morphology relative to age-normed trajectories of brain structural development. Centile Interpretation: Scores near 0.5 reflect typical morphological development for a given age group.

No significant linear or non-linear relationship was found between anatomical centile scores and participant differentiability: r = 0.08, p = 0.11 (Figure S6c).

Overall, these analyses suggest that while anatomical features exhibit age-related changes in similarity patterns, they do not meaningfully contribute to participant differentiation as measured by neurophysiological traits. The lack of a significant relationship between anatomical centile scores and differentiability further supports the independence of neurophysiological traits from structural morphology, underscoring their unique value in characterizing individuality across the lifespan.

**Figure S6.**
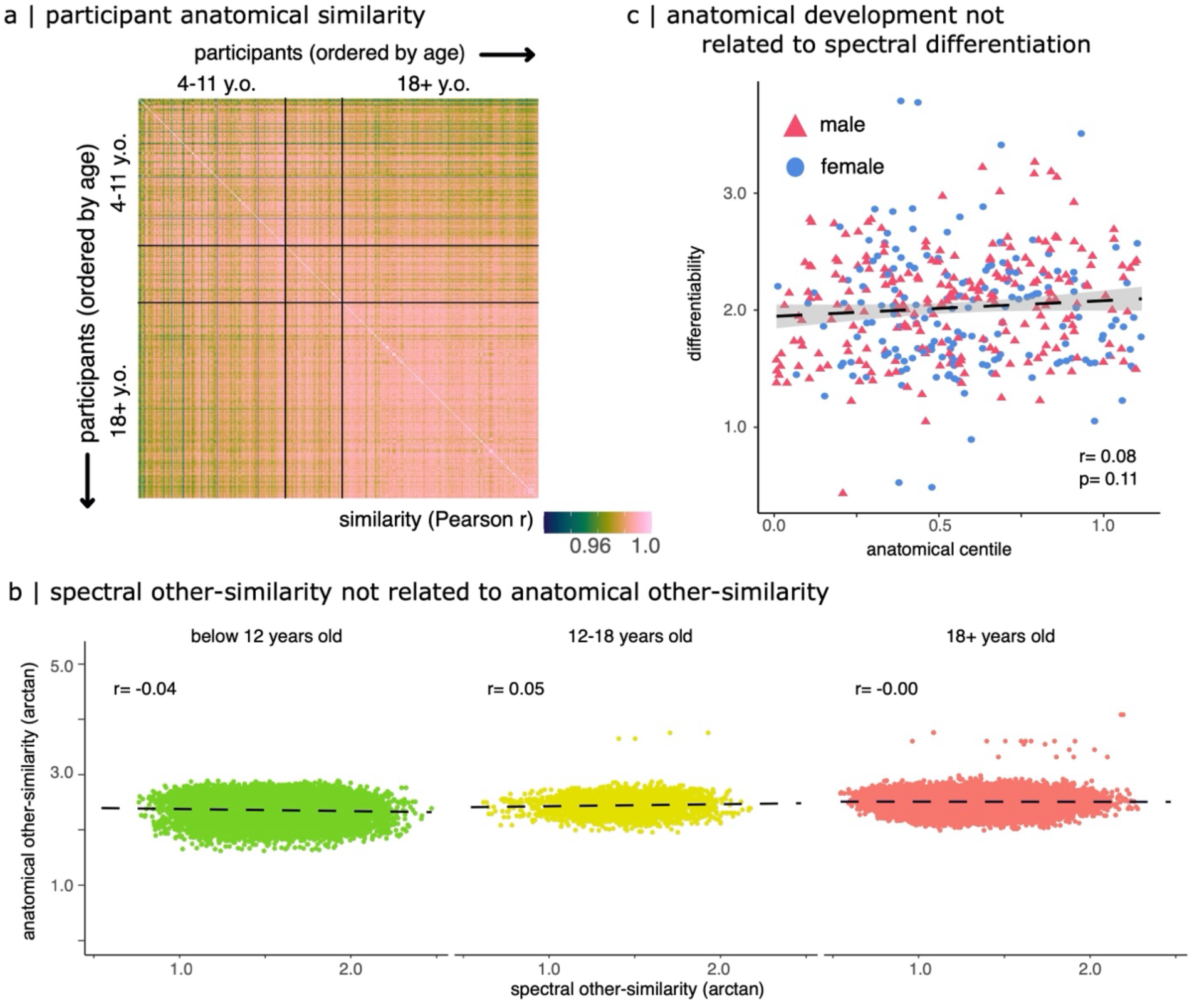
Weak Influence of Anatomy on Differentiation from Neurophysiological Traits. <u>(a) Anatomical Similarity Matrix:</u> The similarity matrix, based on cortical FreeSurfer features (e.g., cortical thickness, surface area, and curvature), orders participants by ascending age. Unlike neurophysiological traits, anatomical features of children do not exhibit greater similarity to other children, as seen in the off-diagonal elements. <u>(b) Lack of Linear Relationship Between Anatomical and Neurophysiological Similarity:</u> The other-similarity of wideband neurophysiological traits (off-diagonal elements from Figure 1a) does not scale linearly with anatomical similarity (off-diagonal elements from panel a). This finding holds across all age groups, indicating weak anatomical influence on neurophysiological differentiation. <u>(c) Role of Anatomical Centile Scores in Differentiation:</u> The anatomical centile score (x-axis) represents an age-normed morphological benchmark, with values near 0.5 indicating average development for a given age group. Participants with extreme centile scores (e.g., atypical morphology) are not easier to differentiate, suggesting anatomical variability does not significantly impact participant differentiation.

### Robustness to age-group definition

To assess the robustness of our differentiation accuracy results, we examined whether changes in the definition of age groups influenced our findings. Specifically, we redefined the age groups as follows:

- **Children**: Participants below 11 years old,
- **Adolescents**: Participants between 11 and 20 years old,
- **Adults**: Participants aged 20 years and older.

Using full spectral features, we observed:

- **Children** (**below 11 years):** Differentiation accuracy **of 89.9%** (CI [88.2, 91.6]),
- **Adolescents (11–20 years):** Differentiation accuracy of **96.9%** (CI [95.7, 98.6]),
- **Adults (20 years and above):** Differentiation accuracy of **95.4%** (CI [94.1, 97.1]).

These results are consistent with our original findings (Figure S7) and demonstrate that the observed differentiation patterns are robust to variations in age-group definitions.

The robustness of differentiation accuracy across alternative age-group definitions underscores the stability of neurophysiological traits as reliable markers of individuality. This consistency further supports the validity of our approach in capturing lifespan changes in neurophysiological traits.

**Figure S7.**
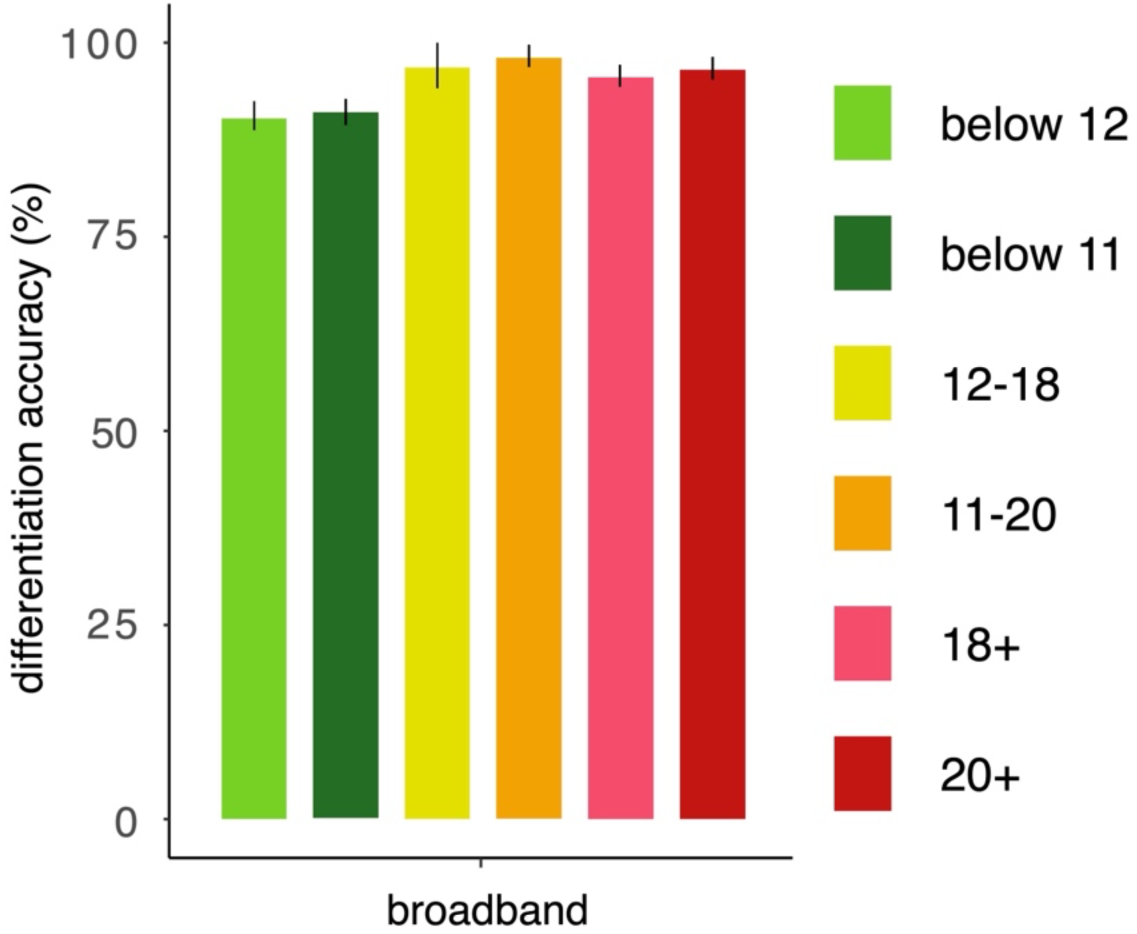
Differentiation Accuracy is Robust Across Age-Group Definitions. Differentiation accuracy remains consistent regardless of the age-group definitions used. Children below 11 or 12 years of age exhibit lower differentiation accuracy compared to adolescents and adults, indicating that younger individuals are harder to differentiate from one another based on wideband neurophysiological traits. The differentiation accuracy for children is consistently lower than that of adults, demonstrating the robustness of the findings to age-group thresholds. Error bars represent bootstrapped 95% confidence intervals.

### Differentiation from Aperiodic and Periodic Traits

To investigate the contributions of aperiodic and periodic brain activity to participant differentiation, we performed analyses as described in the Methods section.

In the Cam-CAN dataset, we observed that aperiodic components supported accurate participant differentiation:

- Entire cohort: 92.6% accuracy (CI [90.9, 94.2])
- Young adults (18–45 years): 91.7% accuracy (CI [88.6, 94.9])
- Middle-aged adults (45–65 years): 95.4% accuracy (CI [93.7, 97.1])
- Older adults (65–89 years): 94.4% accuracy (CI [92.6, 96.0])

Participant differentiation from periodic components showed comparable performance to broadband results:

- Entire cohort: 96.6% accuracy (CI [95.1, 98.1])
- Young adults (18–45 years): 96.7% accuracy (CI [93.7, 99.4])
- Middle-aged adults (45–65 years): 97.3% accuracy (CI [96.6, 98.3])
- Older adults (65–89 years): 97.6% accuracy (CI [96.6, 98.8])

Overall, these results highlight that while aperiodic activity reliably supports participant differentiation, periodic components offer consistently higher accuracy. This underscores the unique role of periodic activity in encoding individualized neurophysiological traits across all age groups.

**Figure S8.**
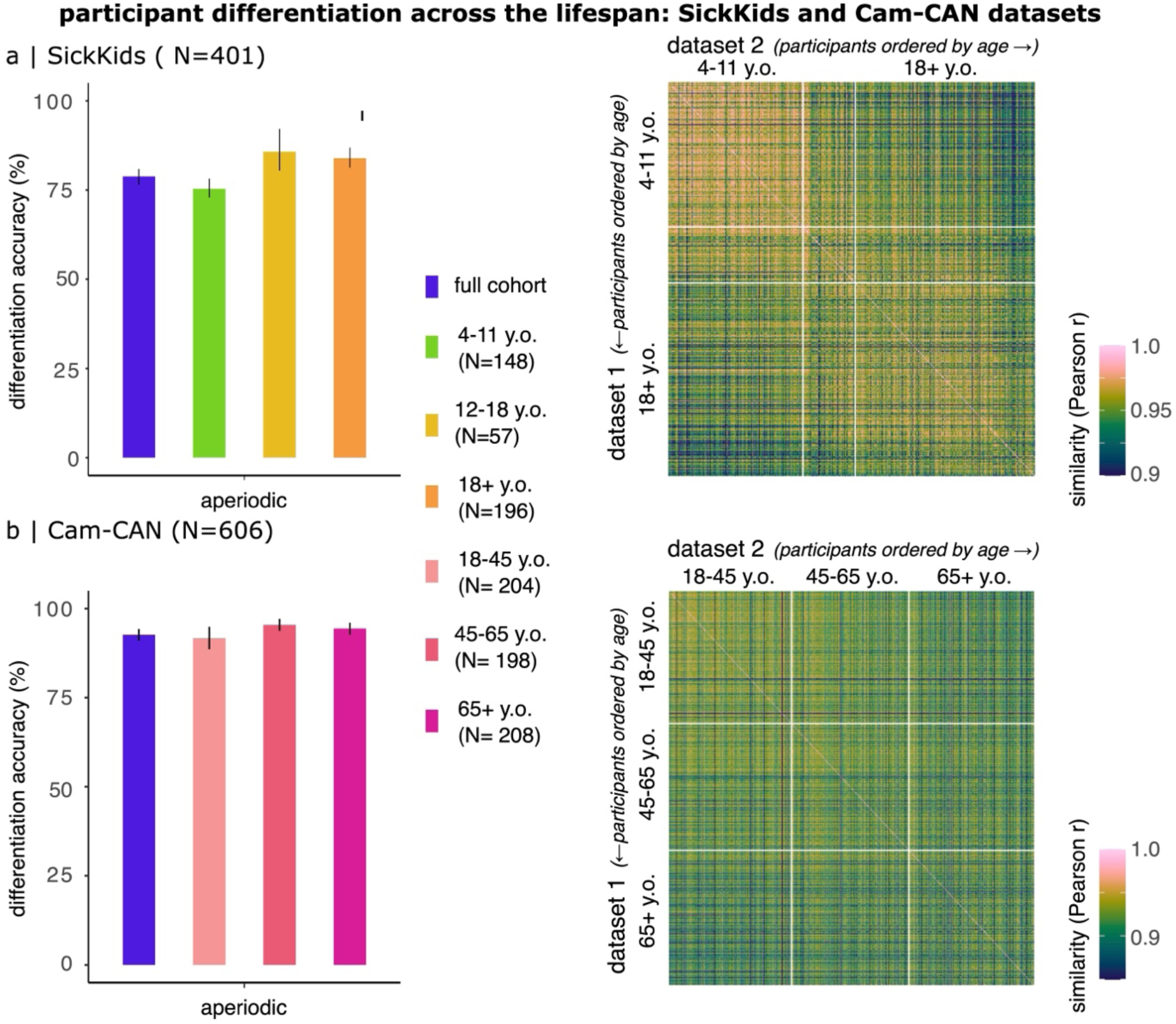
Differentiation from Aperiodic Neurophysiological Traits. Left Panels: Accuracy in distinguishing participants based on aperiodic neurophysiological traits in the SickKids cohort (a) and the CamCAN cohort (b). Error bars represent bootstrapped 95% confidence intervals. Differentiation accuracy is lower for children compared to adults, indicating greater similarity among children’s aperiodic traits. Right Panels: Similarity matrices for aperiodic neurophysiological traits across two data segments, ordered by ascending age.

- SickKids cohort (a): The aperiodic traits of children are more similar to those of other children, as shown by higher values in the off-diagonal elements of the similarity matrix. This increased other-similarity contributes to the challenges in differentiating children based on aperiodic brain activity.
- CamCAN cohort (b): No such effect is observed in adults, as aperiodic neurophysiological traits display lower other-similarity, supporting higher differentiation accuracy in older age groups.

**Figure S9.**
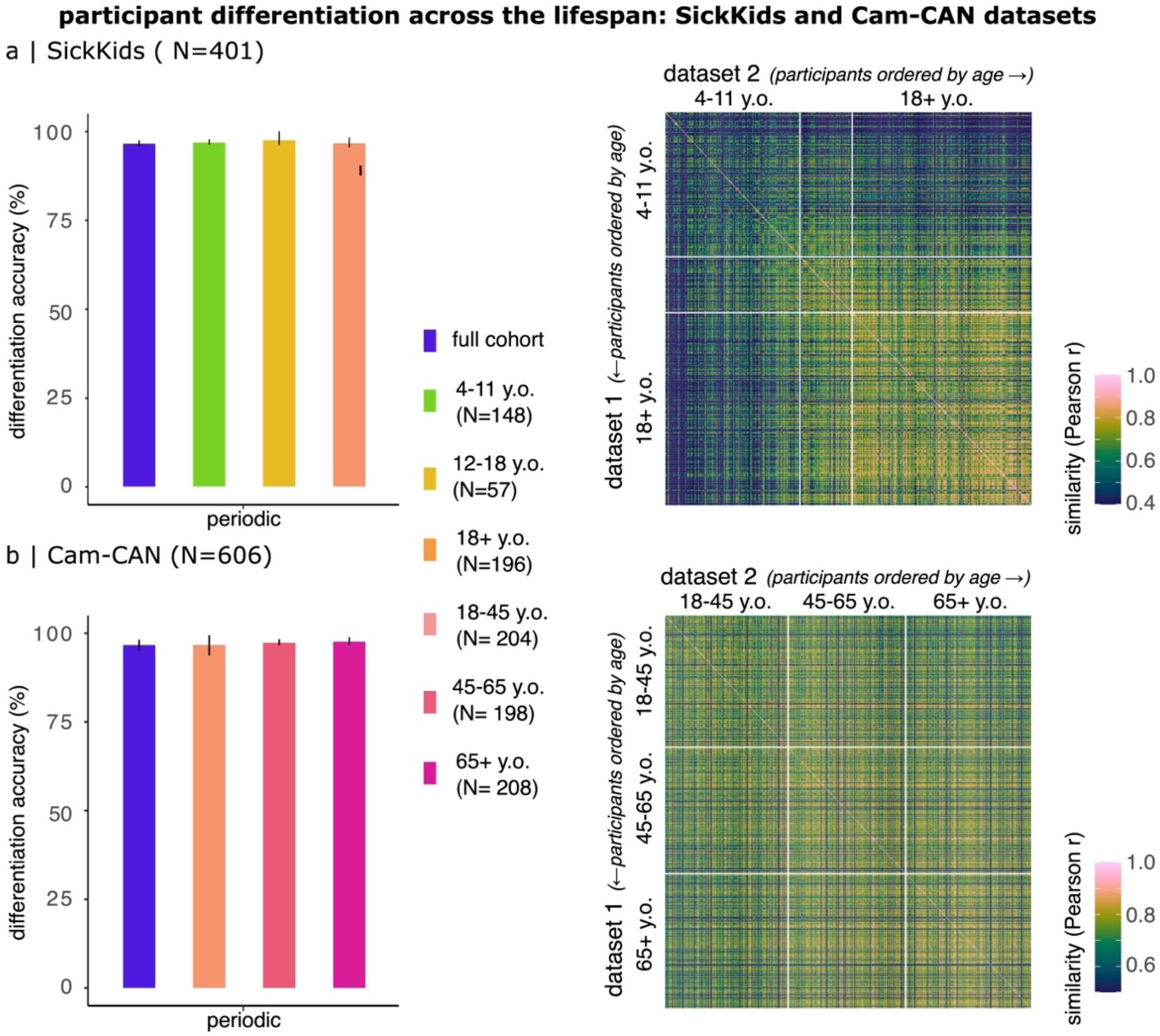
Differentiation from Periodic Neurophysiological Traits. Left Panels: Accuracy in distinguishing participants based on their periodic neurophysiological traits in the SickKids cohort (a) and the CamCAN cohort (b). Error bars represent bootstrapped 95% confidence intervals. Differentiation accuracy remains high across all age groups, reflecting the robustness of periodic brain activity as a marker of individuality. Right Panels: Similarity matrices for periodic neurophysiological traits across two data segments, ordered by ascending age.

- SickKids cohort (a): Participant differentiability from periodic neurophysiological traits shows a weak positive relationship with age, indicating slightly greater distinctiveness in older children compared to younger ones.
- CamCAN cohort (b): Differentiability stabilizes in adulthood and remains consistent into late adulthood, as shown by relatively uniform off-diagonal values across age groups in the similarity matrix.

### Decoding of Cognition from Periodic Neurophysiological Traits

Based on prior fMRI research^11^, we hypothesized that spectral neurophysiological traits, whichdifferentiate individuals, could also predict fluid intelligence scores as measured by the Cattell test^62,103,104^. To test this, we decoded fluid intelligence scores from participants’ periodic neurophysiological traits in the Cam-CAN cohort.

To reduce the high dimensionality of the neurophysiological data, we first applied a principal component analysis (PCA), retaining 148 components that explained 100% of the variance in the periodic neurophysiological traits. We trained Support Vector Regressions (SVR) models on PCA-reduced features using default parameters in R. For model training and validation, we used an 80/20 split, training the model on 80% of the participants and testing on a held-out 20%. This process was repeated 1,000 iterations to ensure robustness.

Our analysis demonstrated that fluid intelligence scores could be reliably decoded from periodic neurophysiological traits, with a mean correlation between decoded and observed scores across iterations of r = 0.52 (t = 14.9, p < 0.001; Figure S10). We also observed a strong correlation between chronological age and fluid intelligence: observed fluid intelligence and age: r = -0.67 (t = -22.1, p < 0.001); decoded fluid intelligence and age: r = -0.68 (t = -22.4, p < 0.001).

These results indicate that resting-state periodic brain activity encodes meaningful cognitive information, such as fluid intelligence, beyond its role in individual differentiation. The correlation between age and fluid intelligence further supports the developmental relevance of these periodic neurophysiological traits.

**Figure S10.**
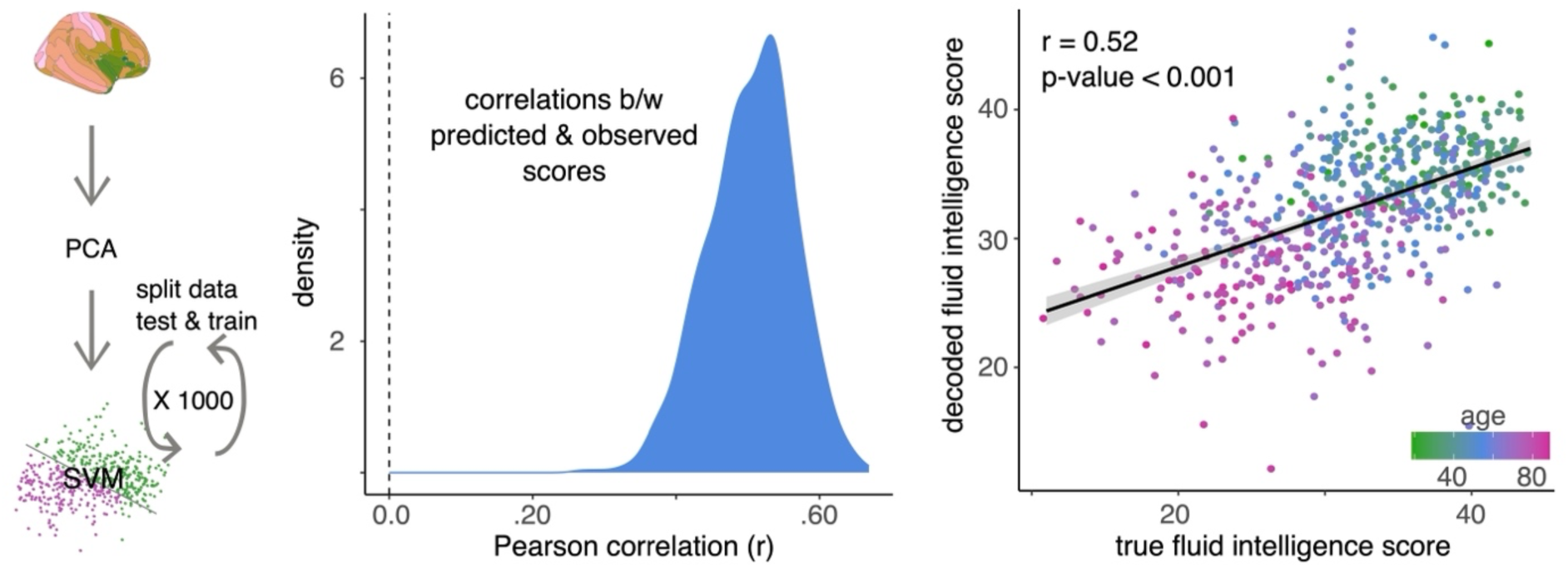
Decoding of Fluid Intelligence Scores from Periodic Neurophysiological Traits in Adults. Left Panel: Pipeline for training Support Vector Regression (SVR) models to decode fluid intelligence performance (Cattell test scores) using periodic neurophysiological traits reduced using Principal Component Analysis (PCA) before model training. Middle Panel: Histogram showing the Pearson correlation coefficients between observed and predicted fluid intelligence scores across 1000 iterations of the SVR model. The high mean correlation indicates reliable decoding of fluid intelligence scores from periodic neurophysiological traits. Right Panel: Scatter plot of decoded fluid intelligence scores (y-axis) versus observed fluid intelligence scores (x-axis). Points are coloured by participants’ chronological age. The strong linear relationship highlights the predictive power of periodic neurophysiological traits for cognitive performance.

**Figure S11.**
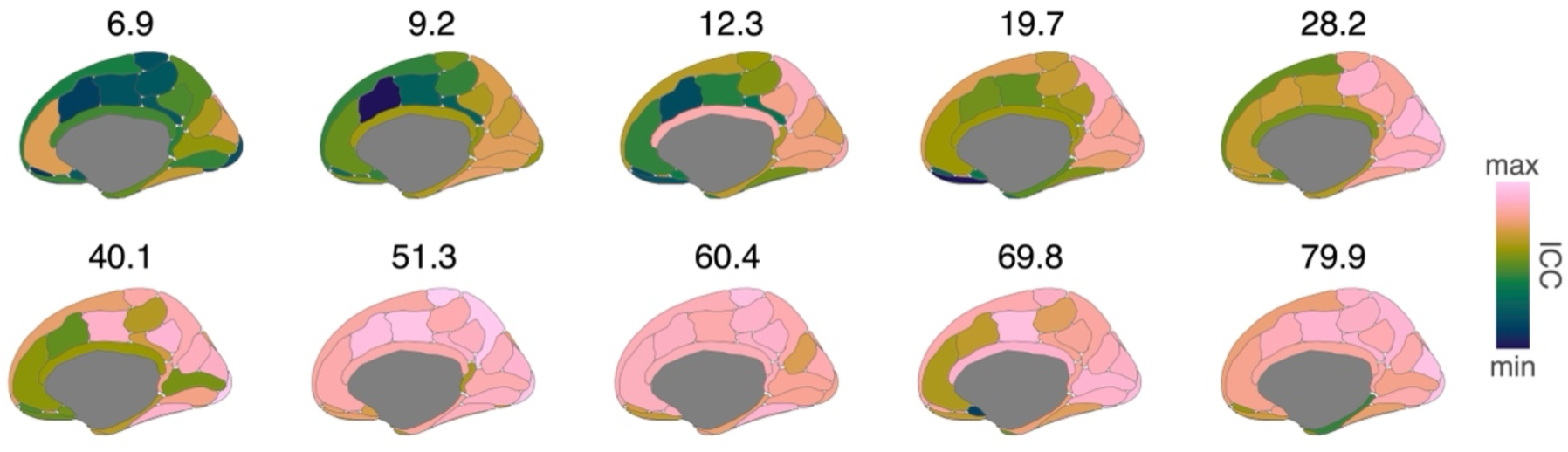
Distinctive Cortical Regions for Differentiation Across the Lifespan: Medial View. Medial views display Intraclass Correlation Coefficients (ICCs) for cortical regions contributing most to individual differentiation of neurophysiological activity across different age groups (bins of 100 individuals). Higher ICC values indicate regions with greater differentiation across individuals. These medial views complement the lateral maps shown in Figure 2a, providing a comprehensive depiction of distinctive regions across the cortex.

**Figure S12.**
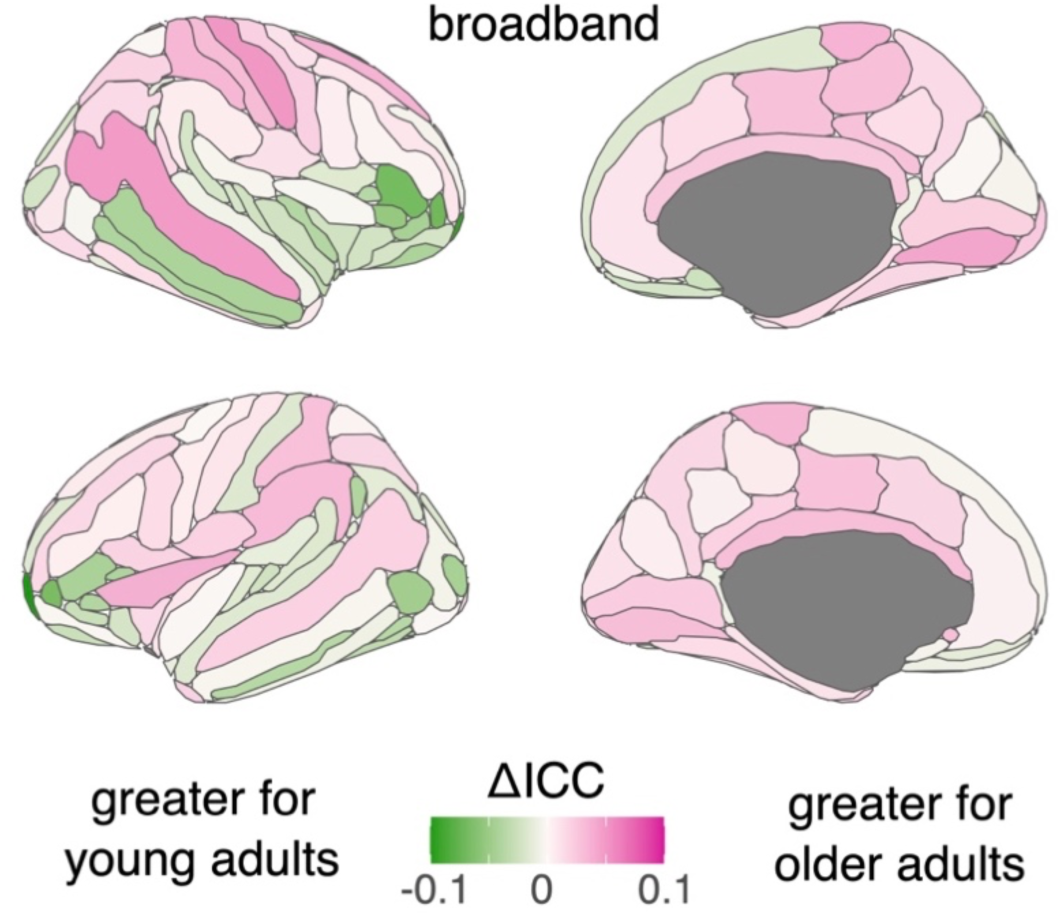
Sensorimotor Regions Are More Characteristic of Older Adults. Maps display the topographic distribution of age-related differences in the saliency of characteristic cortical regions (ΔICC) for wideband neurophysiological traits in the Cam-CAN cohort. Shades of green indicate regions that are more characteristic of young adults (18–45 years old), while sahdes of pink highlight regions that are more characteristic of older adults (65+ years old). The findings underscore the increasing importance of sensorimotor regions in differentiating neurophysiological traits with age.

**Table S6.** The nonlinear effect of age on the alignment between differentiable brain activity and the unimodal-to-transmodal functional gradient.

| arctanh(alignment) |  |  |  |
| --- | --- | --- | --- |
| Predictors | Estimates | CI | p |
| (Intercept) | -0.31 | -0.33 – -0.28 | **<0.001** |
| group [1st degree] | -0.12 | -0.27 – 0.02 | 0.097 |
| group [2nd degree] | 0.24 | 0.09 – 0.39 | **0.003** |
| group [3rd degree] | -0.46 | -0.60 – -0.31 | **<0.001** |
| Observations | 37 |  |  |
| R<sup>2</sup> / R<sup>2</sup> adjusted | 0.614 / 0.579 |  |  |

**Figure S13.**
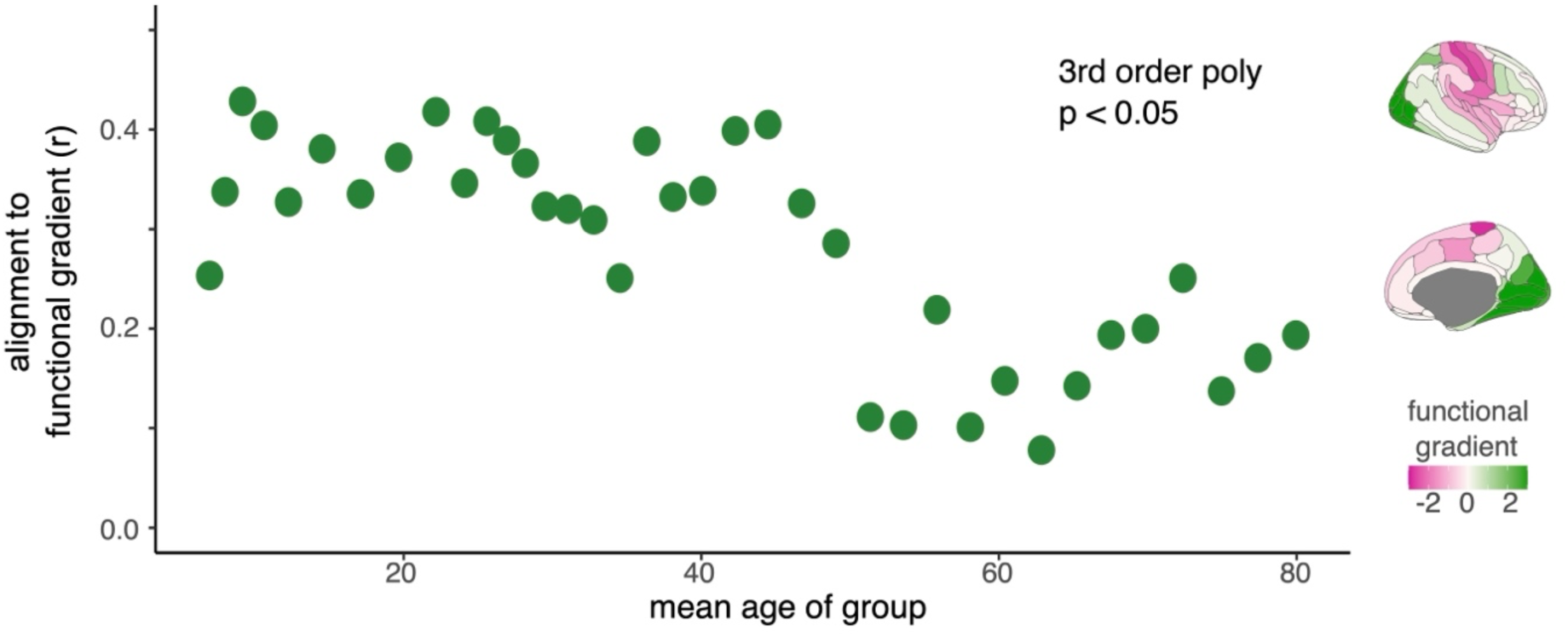
Alignment Between Distinctive Cortical Regions and the Visual-to-Motor Functional Gradient. Scatter plot illustrating the alignment between distinctive cortical regions for differentiation (Figure 2a) and the visual-to-motor functional gradient (topography shown on the right). The alignment varies across age groups, following a non-linear trajectory, though the relationship did not survive correction for spatial autocorrelation. The strongest alignment was observed in childhood (9.2 years old), while the weakest alignment occurred in late adulthood (62.8 years old). These findings highlight the dynamic relationship between neurophysiological differentiation and functional gradients across the lifespan.

**Figure S14.**
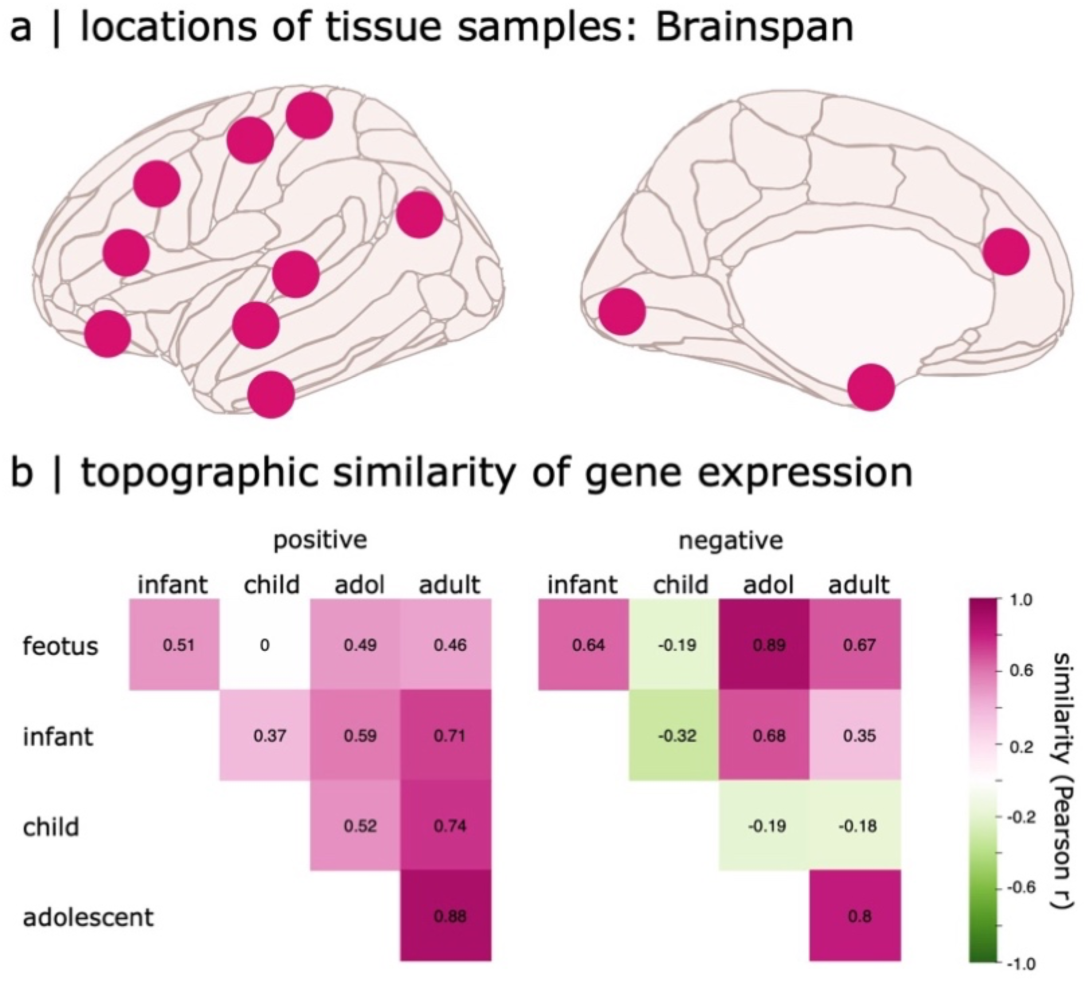
The Positive Gene Signature is Topographically Stable. (a) Approximate cortical sampling locations of the BrainSpan dataset^98^, spanning multiple neurodevelopmental stages. (b) Topographic similarity of gene expression profiles for the identified positive (right panel) and negative (left panel) gene signatures across five developmental stages^70,98,99^: fetus (8–37 post-conception weeks), infant (4 months–1 year), child (2–8 years), adolescent (11–19 years), and adult (21–40 years). The positive gene signature remains highly stable across these stages (Pearson r > 0.7), suggesting a conserved spatial pattern of gene expression related to participant differentiation throughout neurodevelopment.

**Table S7.** Alignment with the Positive Gene Signature.

| arctanh(alignment) |  |  |  |
| --- | --- | --- | --- |
| Predictors | Estimates | CI | p |
| (Intercept) | 0.46 | 0.42 – 0.49 | **<0.001** |
| group [1st degree] | -0.46 | -0.69 – -0.24 | **<0.001** |
| group [2nd degree] | -0.27 | -0.50 – -0.05 | **0.019** |
| group [3rd degree] | 0.73 | 0.51 – 0.96 | **<0.001** |
| Observations | 37 |  |  |
| R<sup>2</sup> / R<sup>2</sup> adjusted | 0.672 / 0.642 |  |  |

